# Does transcriptome of freshly hatched fish larvae describe past or predict future developmental trajectory?

**DOI:** 10.1101/2023.10.27.564361

**Authors:** Rossella Debernardis, Katarzyna Palińska-Żarska, Sylwia Judycka, Abhipsa Panda, Sylwia Jarmołowicz, Jan P. Jastrzębski, Tainá Rocha de Almeida, Maciej Błażejewski, Piotr Hliwa, Sławomir Krejszeff, Daniel Żarski

## Abstract

Transcriptomic analysis of freshly hatched fish larvae offers insights into phenotype development, yet it remains uncertain whether it reflects parental influence or predicts individual growth. This research scrutinizes the transcriptome of 16 Eurasian perch (*Perca fluviatilis*) larval groups alongside pre- and post-hatching traits. Despite consistent fertilization rates, significant variations in embryonic development and larval performance highlight diverse phenotypes studied. It enabled to bring our attention to the fact that larval transcriptome can serve as a window into both the parental contributions and the future performance of the larvae. Our further analysis shed light on ribosome biogenesis, neurogenesis, and the cell cycle, as important processes shaping early larval performance. Additionally, we propose a set of predictive, validated gene markers associated with further larval performance and key aquaculture traits, such as *selenoo* (associated with cannibalism), *trim16* (fulton’s condition factor), *slc15a1* (specific growth factor), and *cipc* (final weight). This study delves into the descriptive and predictive nature of the transcriptomic portrait of newly hatched larvae, paving the way to comprehend the intricate developmental pathways from fertilization towards juvenile stage.

## 1. Introduction

The early life history of most teleost fishes (both marine and freshwater) embraces a crucial stage known as larval period (McMenamin and Parichy, 2013). This transitional phase involves changes across morphological, behavioral, and physiological levels which lead to individual’s progress towards adulthood (Urho, 2002). The dynamics and vulnerability of this phase are what makes larvae important to study in order to facilitate a better understanding of the developmental journey from the egg stage to subsequent juvenile and adult stages.

Larval phase is characterized by significant events encompassing the initiation of the exogenous feeding, yolk-sac reduction, and the inflation of the swim bladder, as well as the development and functional maturation of different organs, tissues and systems (e.g.: nervous, visual, digestive and immune systems) (Osse et al., 1997). These changes are regulated by a cascade of events as well as intrinsic (e.g. genetics, physiology) and extrinsic factors (e.g., temperature, density, nutrition (Sarropoulou et al., 2016)) which have been a longstanding focus for the scientific community. Therefore, due to their significant influence on later developmental phases, investigations on the factors and intrinsic processes influencing larval performance are of high importance. This could additionally directly impact the modification of breeding and hatchery protocols by aquaculturists.

Sustainable development of aquaculture industry requires species diversification and the intensification of production to meet the rising global demand for aquatic products (Subasinghe et al., 2009). Among cultured species, Eurasian perch (*Perca fluviatilis*) is considered a valuable model due to its commercial relevance and practise in intensive farming using recirculating aquaculture systems (RAS). Although, its production is already well-established (Fontaine and Teletchea, 2019; Palińska-Żarska et al., 2020; Policar et al., 2019), variable reproductive and larviculture performances remain still an obstacle to the improvement of its breeding efficiency. Within this framework, the production of high-quality larvae becomes paramount, as they possess a significant capacity to adapt to the aquaculture environment (Koumoundouros et al., 2017; Valente et al., 2013). Despite substantial progress in comprehending larvae biology and establishing larviculture protocols for many fish species, achieving consistently high survival rates and optimal growth potential remains a challenge (Valente et al., 2013). To overcome this, a deeper understanding of intrinsic and extrinsic factors influencing larval environmental adaptability is required. This is crucial for establishing a clear definition or, at the very least, identifying descriptors or predictors of larval quality, addressing an urgent need in the field.

Currently, the assessment of fish larvae quality is predominantly tied to morphological traits, such as body shape (i.e., absence of skeletal deformities), yolk sac morphology, inflated swim bladder, pigmentation (i.e., Koumoundouros, 2010; Boglione et al., 2013; Koumoundouros et al., 2017). Despite their significance, these indicators have limitations. Assessing these traits requires laborious and long-term breeding operations and their accuracy may be compromised by potential biases resulting from the rearing environment and human interventions. For this reason, there is a growing emphasis on incorporating new molecular quality indicators (e.g., gene markers) for larvae to supplement existing methods. The integration of molecular approach with conventional morphological examination will improve the efficiency of larval rearing protocols by offering a comprehensive understanding of biological processes. These molecular signatures should be found considering larvae at the earliest stage (preferably after hatching), where individuals are fully autonomous but major human intervention which could cause some stress and thus affect the overall regulation of biological processes of the fish is not yet in place (Valente et al., 2013). For Eurasian perch larvae, the mouth opening stage represents this time point (Palińska-Żarska et al., 2021). However, despite multiple evidences suggesting the potential of larval molecular profiles in predicting performance, there is still a lack of specific molecular indicators or fingerprints as reliable indicators. This stems from the lack of studies that directly align the characterisation of molecular profiles against larval traits.

Transcriptomics has been already successfully applied to fish larvae biology research, offering valuable insights into the molecular mechanisms underlying various biological processes (Chandhini and Rejish Kumar, 2019; Ferraresso et al., 2013; Mazurais et al., 2011; Żarski et al., 2017e). While it underscores the significance of larval transcripts as a proxy to understand organismal phenotypes, it remains unclear if such profiles can describe/predict larval outcomes. Therefore, the current study on Eurasian perch aims to explore whether the transcriptomic profile of 16 different groups of larvae at the mouth opening stage, combined to zootechnical features, could be an indicative tool for describing the past and predicting the future fate of the larvae by highlighting gene markers or molecular mechanisms that could serve as indicators of larval performance-related traits.

## 2. Material and Methods

Sixteen diverse groups of Eurasian perch larvae were obtained by controlled reproduction of wild spawners from different water bodies (Supplementary file S1). A simple mating design was carried out by crossing a single female with an individual male.

### 2.1. Broodstock management and origin

#### 2.1.1. Males’ origin and management

The Eurasian perch males (average weight 190.5 ± 58 g), used for controlled reproduction, were from earthen pond systems located in Central (Rytwiany and Łyszkowice Fish Farm) and North of Poland (Iława Fish Farm). Spawners were harvested during late autumn (end of October and early November) and overwintered in the flow-through system of the Salmonid Research station of the National Inland Fisheries Research Institute (IRS-PIB) in Rutki (North Poland) under the natural photoperiod. Male individuals were captured in autumn since it is very difficult to catch them during the breeding season, and moreover, often completed or already contributed to the spawning act before being caught, which may affect sperm quality. Then, during the spawning period, males were transferred in plastic bags with oxygen (Żarski et al., 2017) to the Center of Aquaculture and Ecological Engineering of the University of Warmia and Mazury in Olsztyn (CAEE-UWM, NE Poland), where they were placed in the RAS with a controlled photoperiod (14 L:10 D) and temperature (12°C) until spermiation took place. The sexually mature fish were hormonally stimulated (with an intraperitoneal injection at the base of the left ventral fin) using a salmon gonadoliberin analog (sGnRHa, Bachem Chemicals, Switzerland) at a dose of 50 μg kg−1(Żarski et al., 2020). Before any manipulation, individuals were anesthetized in MS-222 (Argent, USA) at a concentration of 150 mg l^−1^. Total length (TL), fork length (FL) and body weight (before stripping) were measured for each individual (Supplementary file S1).

#### 2.1.2. Sperm sampling and cryopreservation protocols

Five days after hormonal injection, semen was collected using a catheter (to avoid contamination of the urine;(Sarosiek et al., 2016)) and with gentle abdominal pressure (i.e., stripping). After collection, each sample was kept on ice, and sperm motility parameters were evaluated with CASA system (Supplementary file S2), (using the CEROS II system -Hamilton-Thorne, USA-; as described by Judycka et al. (2022), while the sperm concentration of fresh semen was measured using a NucleoCounter SP-100 computer aided fluorescence microscope (Chemometec, Allerød, Denmark;(Nynca and Ciereszko, 2009)). For this purpose, the semen was first diluted 100 times with PBS and then 51 times with Reagent S100 and loaded into the kit cassette containing propidium iodide. SemenView software (Chemometec, Denmark) was used to determine the final concentration of spermatozoa in each sample (Judycka et al., 2019).

Thereafter, sperm cryopreservation was carried out following the procedure described by Judycka et al. (2022). Briefly, the semen was diluted with a glucose-methanol (GM) extender supplemented with potassium chloride (consisting of a final concentration of 0.30 M glucose, 7.5% methanol and 25 mM KCl at 3.0 × 109/ml spermatozoa). Semen mixed with cryoprotectants was filled into 0.5 ml plastic straws and then placed on a floating rack and cryopreserved in liquid nitrogen vapor for 5 min. Next, the straws were submerged into liquid nitrogen, which ended the process. After cryopreservation, the sperm motility was re-evaluated by thawing the straws in a water bath for 10s at 40°C. Finally, the straws were placed in liquid nitrogen storage tanks until being used for fertilization. For this operation, cryopreserved semen was used to ensure feasibility of the entire operation (in case of delayed reproduction of females) and to maintain comparable sperm quality in case the females did not ovulate simultaneously (collection of eggs from wild females during the spawning season can take several days; (Żarski et al., 2017b)) and is currently used as a standard procedure in many selective breeding programs (Judycka et al., 2022).

#### 2.1.3. Females’ origin and management

For this experiment, 16 wild females (average weight 493 ± 213 g) were utilized. Some individuals came from the Żurawia and Iława pond systems, which were overwintered together with males (as mentioned before). In addition, in April, wild females from Szymon and Umląg Lakes were captured using gill nets during the spawning season. In this way, we ensured significant phenotypic variability, expected to provide appropriate heterogeneity in the performance of the larvae, which was needed to fulfil the purpose of this research project. Prior to spawning, all fish were transported in plastic bags with oxygen to the CAEE-UWM where they were placed in the RAS with a controlled photoperiod (14 L:10 D) and temperature (12°C) until ovulation. Prior to hormonal stimulation, females were first catheterized, and the oocyte maturation stage was determined following the classification proposed by Żarski et al. (2011). After that, the fish were hormonally stimulated (as with males) using a salmon gonadoliberin analog (sGnRHa, Bachem Chemicals, Switzerland) at a dose of 50 μg kg^−1^ (Żarski et al., 2020). Before any manipulation, individuals were anesthetized in MS-222 (Argent, USA) at a concentration of 150 mg l^−1^. During the experiment, for each fish, the total length (TL), fork length (FL), body weight and weight of the ribbon were taken (measures are provided in Supplementary file S1).

#### 2.1.4. Egg collection and fertilization protocols

At ovulation, the eggs (ribbon) were collected through hand stripping (gentle massage of the abdomen part of the fish body as described by Żarski et al. (2011)). After stripping, the number of dry eggs in 1 g was evaluated by first counting the eggs in 3 small portions (∼0.2 g each) of the ribbon (Żarski et al., 2017b). In this way, the correct number of spermatozoa to be used for fertilization of each ribbon was estimated (Żarski et al., 2017c). A ribbon with an average weight of 80 ± 12 g was used to carry out the *in vitro* fertilization as described by Judycka et al. (2019). Briefly, the eggs were first activated with modified Lahnsteiner activating solution (75 mM NaCl, 2 mM KCl, 1 mM MgSO_4_×7H_2_O, 1 mM CaCl_2_× 2H_2_O, 20 mM Tris, pH 8 (Judycka et al., 2022). Hereafter, just before fertilization, straws with cryopreserved semen were thawed in a water bath at 40°C for 10 s and placed in an Eppendorf tube. Thirty seconds after egg activation, sperm was added at a sperm:egg ratio of 200 000:1. The eggs were then stirred for 40 seconds and washed with hatchery water after ∼10 minutes to remove excess sperm and any debris. The same procedure was followed for each female separately.

#### 2.1.5. Egg incubation and hatching

The fertilized eggs were incubated in 15 L black-walled tanks (eggs from each female separately) operating in the same RAS and placed on nets with mesh diameter of 3 mm at a temperature of 14°C. After 12 hours post fertilization (HPF), ∼100 eggs from each batch were randomly sampled (in duplicate) to evaluate the fertilization rate before the maternal-to-zygotic transition (MZT), which occurs at around 13 HPF (Güralp et al., 2016), and then the embryonic development at the neurula stage (when the body of the embryo can already be easily seen, at around 3 days post fertilization). During egg incubation, the temperature was raised to 15°C when the embryos reached the eyed-egg stage and then to 16°C as soon as the first spontaneously hatched larvae were observed. In Eurasian perch, hatching can last for 5 days, even for the same batch of eggs (Żarski et al., 2017d). Therefore, to ensure almost synchronous hatching, manual hatching was induced. Briefly, the eggs were moved to a bowl (each batch separately) and stirred gently. The hatched larvae were then moved back into the 15L tanks. This operation was repeated several times until most of the larvae hatched. This moment was considered the end of hatching (0 days post hatching - DPH).

After hatching, the larvae were left for 24 h without any human interaction, and next, all the larvae within the groups were volumetrically counted and stocked back to the rearing tanks with the same stocking density for each tank and each group (∼ 2500 larvae per tank). Larvae from each batch were stocked into 3 separate tanks constituting separate replicates.

#### 2.1.6. Larviculture and advanced zootechnics

Every group of Eurasian perch larvae was reared in triplicate in the same RAS conditions, following a set of validated and standardized larval rearing methods (i.e., advanced zootechnics) as described by Palińska-Żarska et al. (2020). The larvae were exposed to a specific thermal regime, photoperiod, and feeding schedule (Fig. 1A). The water temperature was automatically controlled throughout the rearing period. After hatching, at 0 DPH, the temperature was 16°C. At 1 DPH, the water temperature was raised by 1°C, and at 2 DPH, it was at 18°C, which was kept stable up to 10 DPH. From 11 DPH onward, the water temperature was gradually increased by 1°C per day until 23°C, considered the optimal temperature for the growth of Eurasian perch larvae (Kestemont et al., 2003; Palińska-Żarska et al., 2020). Starting from 4 DPH, the larvae began to be fed with *Artemia* sp. nauplii *ad libitum* three times per day (first four days of feeding – micro Artemia cysts [SF origin], then standard size Artemia cysts at 260,000 nauplii per gram [GSL origin]) until weaning. From 17 DPH, the larvae were sharply weaned and then fed exclusively dry feed (Perla Larva Proactive, Skretting, Norway) six times a day, pouring it into each tank in small doses, with intervals of 3 minutes, for approximately 15 minutes. At ∼ 30 DPH, the experiment was completed, as with the temperature regime used, and the larval period was considered finished. During the rearing trial, the photoperiod was 24 L:0 D, and the light intensity measured at the water surface was 1500 lux. In addition, the oxygen level and ammonia concentration in the tanks were analyzed every two days. No oxygen level below 80% was ever observed, and the ammonia level was always lower than 0.02 mg L^−1^. The tanks were cleaned twice a day (in the morning, 1 h after feeding, and in the evening just prior feeding), and the dead larvae were collected and counted under a microscope to assess the survival rate (%) throughout the experimental trial. In addition, from 14 DPH, dead larvae were observed under the microscope to assess the cannibalism rate by recording the number of larvae with damage to the body, especially the tail.

**Fig. 1:**
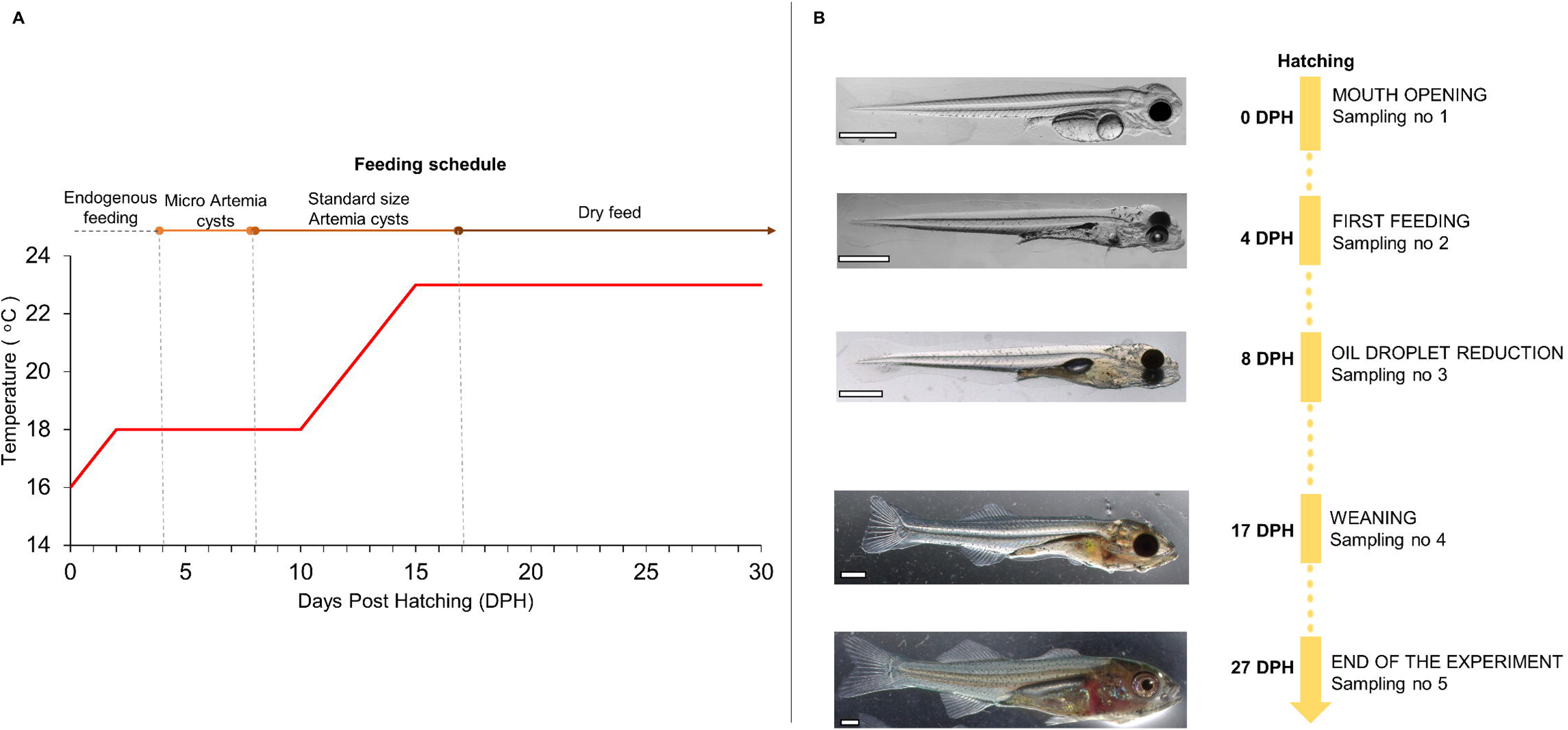
**(A)Temperature regime** (red curve) and the feeding schedule followed during Eurasian perch larvae rearing. **(B) Scheme of Eurasian perch larvae samplings at specific developmental stages**. DPH – Days post hatching, sampling no 1 – moment of mouth opening in at least 50% of larvae, sampling no 2 – moment of exogenous feeding starting in at least 50% of perch larvae, sampling no 3 –oil droplet reduction in at least 50% of larvae, sampling no 4 – time of weaning with dry feed diet, sampling no 5 – end of experiment

### 2.2. Sampling

Larval samplings were carried out at five precise developmental stages (Palińska-Żarska et al., 2021, 2020) (Fig. 1B):

1. At the mouth opening stage (0 DPH),
2. At 4 DPH, when at least 50% of larvae started exogenous feeding,
3. At 8 DPH, in at least 50% of the larvae, almost complete reduction of oil droplets was observed,
4. At the weaning stage (17 DPH),
5. At 27 DPH, considered the end of the experiment, when at least 50% of larvae had undergone the removal of the fin fold.

At each sampling point, 30 larvae per group were sampled to record total length (TL, ±0.01 mm) and wet body weight (WBW, ±0.1 mg). The individuals were first anesthetized (with MS-222, at a concentration of 150 mgL^−1^) and then photographed under a stereoscopic microscope (Leica, Germany) to measure the TL. The WBW was determined with a precision laboratory scale by placing the anesthetized larvae on a nylon net (with a mesh size of approx. 200 μm) and draining the excess water with filter paper (Krejszeff et al., 2013). Additionally, two days after oil droplet reduction (10 DPH), the swim bladder inflation effectiveness (SBIE,%) was evaluated on approximately 100 larvae per experimental group by triple counting (as described by (Palińska-Żarska et al., 2020). Briefly, the individuals were first captured randomly from each tank, placed on a Petri dish, anesthetized and then counted under a stereoscopic microscope (individuals with and without a filled swim bladder). Additionally, to perform molecular analysis, 40 freshly hatched larvae per group were randomly sampled, anesthetized and preserved in RNAlater (Sigma□Aldrich, Germany) following the manufacturer’s recommendations.

### 2.3. RNA extraction

RNA extraction was performed for larvae from each group separately. Using a TotalRNA mini-kit (A&A Biotechnology, Poland), total RNA was extracted from a pool of 10 larvae (n=10, per group), as described by (Palińska-Żarska et al., 2020). Then, the quantity and purity of the extracted RNA were evaluated using a NanoDrop 8000 spectrophotometer (Thermo Fisher Scientific, USA). Samples showed absorbance ratios A260/280 ∼2.0 and A260/230 ∼2.2. The quality of the extracted total RNA was also evaluated using an Agilent Bioanalyzer 2100 (Agilent Technologies, USA), and all the samples presented RIN > 9.0.

### 2.4. RNA-sequencing library preparation

RNA-seq analysis was performed by Macrogen (Amsterdam, Netherlands) using the TruSeq Stranded Total RNA kit (Illumina) with a NovaSeq600 platform, and 40 M 150 bp paired-end reads per sample were generated. The raw reads were quality controlled using FastQC software ver. 0.11.9 (Simon Andrews, 2020). Adapters and low-quality fragments of raw reads (average QPhred score < 20) were trimmed, and reads were clipped to equal lengths of 100 nt using the Trimmomatic tool ver. 0.40 (Bolger et al., 2014). The resulting read sets of the analyzed samples were mapped to a reference genome *P. fluviatilis* obtained from the NCBI database (Sayers et al., 2022) using STAR software ver. 2.7.10a (Dobin et al., 2013) with ENCODE default options. The annotation and estimation of the expression levels were performed using the StringTie tool ver. 2.2.0 (Pertea et al., 2015), with interpreting strand-specific sequencing the ‘fr—firststrand’ parameter was activated. Counts per transcript and gene were calculated using the prepDE Python script (https://github.com/gpertea/stringtie/blob/master/prepDE.py), and TPM and FPKM values were read from StringTie output data using the lncRna R library (Jastrzebski et al., 2023). Functional annotation was performed using eggNOG-mapper (version emapper-2.1.9) (Cantalapiedra et al., 2021) based on eggNOG orthology data (Huerta-Cepas et al., 2019). Sequence searches were performed using DIAMOND (Buchfink et al., 2021).

### 2.5. Construction of the gene co-expression network analysis (WGCNA)

The counts per transcript were employed to perform weighted gene co-expression network analysis (WGCNA) with the R package, following the authors’ recommendations (Langfelder and Horvath, 2008). Briefly, the hierarchical clustering of samples with Euclidean distance was used to check the presence of outliers (Fig 2A). After that, the total gene counts were first filtered by removing all genes with less than 10 counts in each samples. Subsequently, variance-stabilizing transformation was performed, using the DESeq2 package (Love et al., 2014). Then, to proceed with automatic blockwise network construction, the adjacency matrix was calculated, and a soft-power threshold (β=16, leading to signed R^2^ = 0.85, which was the best scale-free indicators for the current analysis) was chosen based on a scale-free topology model (Fig. 2B). Then, a block network signed was constructed and gene modules were identified by hierarchical clustering dendrogram (Fig.3). After, the module eigengene (ME) distances were calculated in order to detect potential relationships of modules with the zootechnical traits collected during the experiment. This will results in gene significance (GS) values and the corresponding p-value for all the modules and traits. To visualize the associations between modules and traits, a module-trait heatmap is generated, offering a graphical representation of the correlation patterns. We focused on the significant modules with a p < 0.05 and the absolute value of the correlation coefficient |cor| ≥ 0.6. Genes embedded in the significant modules of interest were then extracted to proceed with the GO Enrichment analysis.

**Fig. 2:**
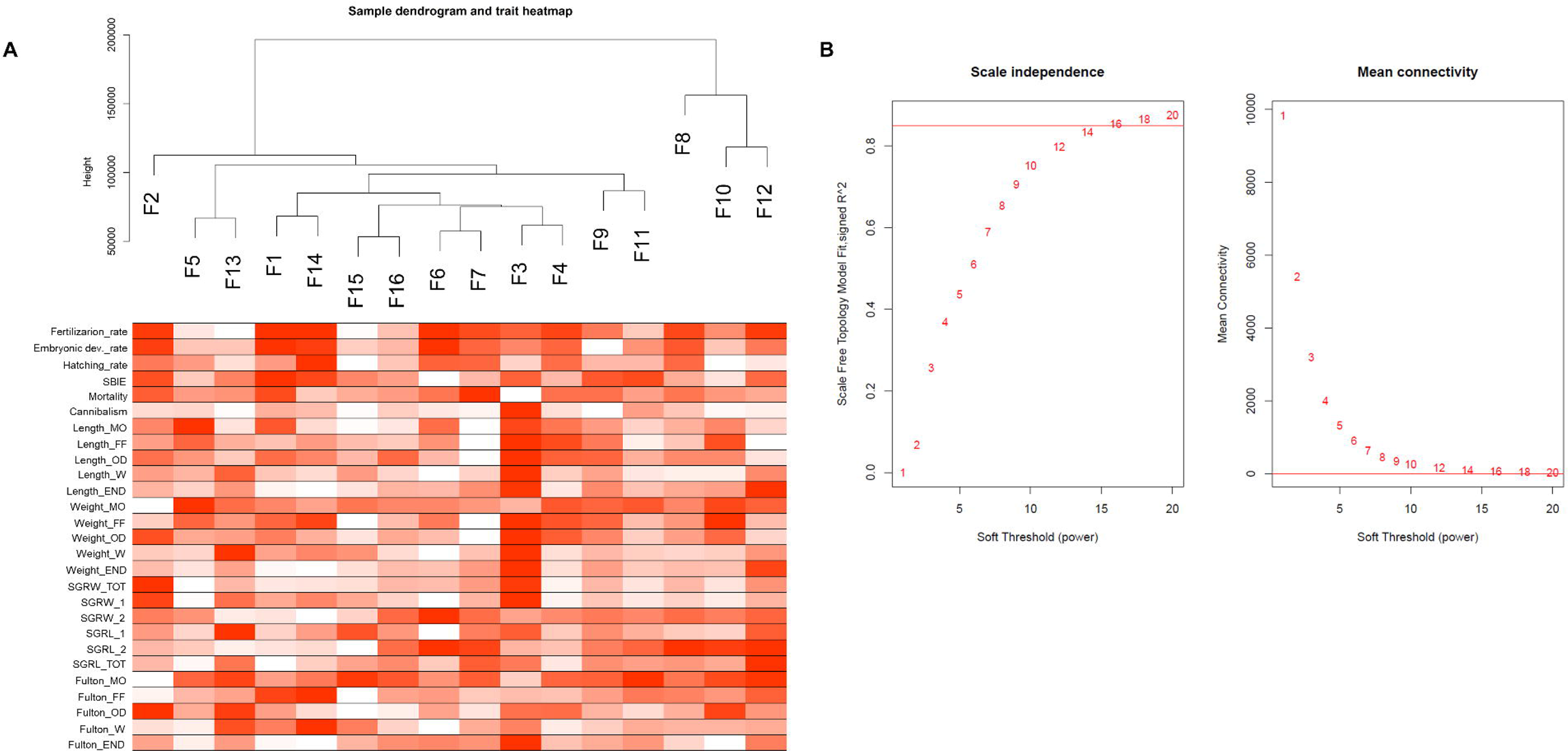
**A. Sample dendrogram based on their Euclidean distance and traits heatmap**. The dendrogram plotted by hierarchical clustering for the 16 experimental groups. The heatmap presented below the dendrogram represents an overview of the zootechnical traits for the corresponding groups. Red colour denotes higher values, while white signifies lower values of traits **B. Determination of soft-threshold power in the WGCNA**. Analysis of the scale-free index for various soft-threshold powers (β) and analysis of the mean connectivity for various soft-threshold powers.

**Fig. 3:**
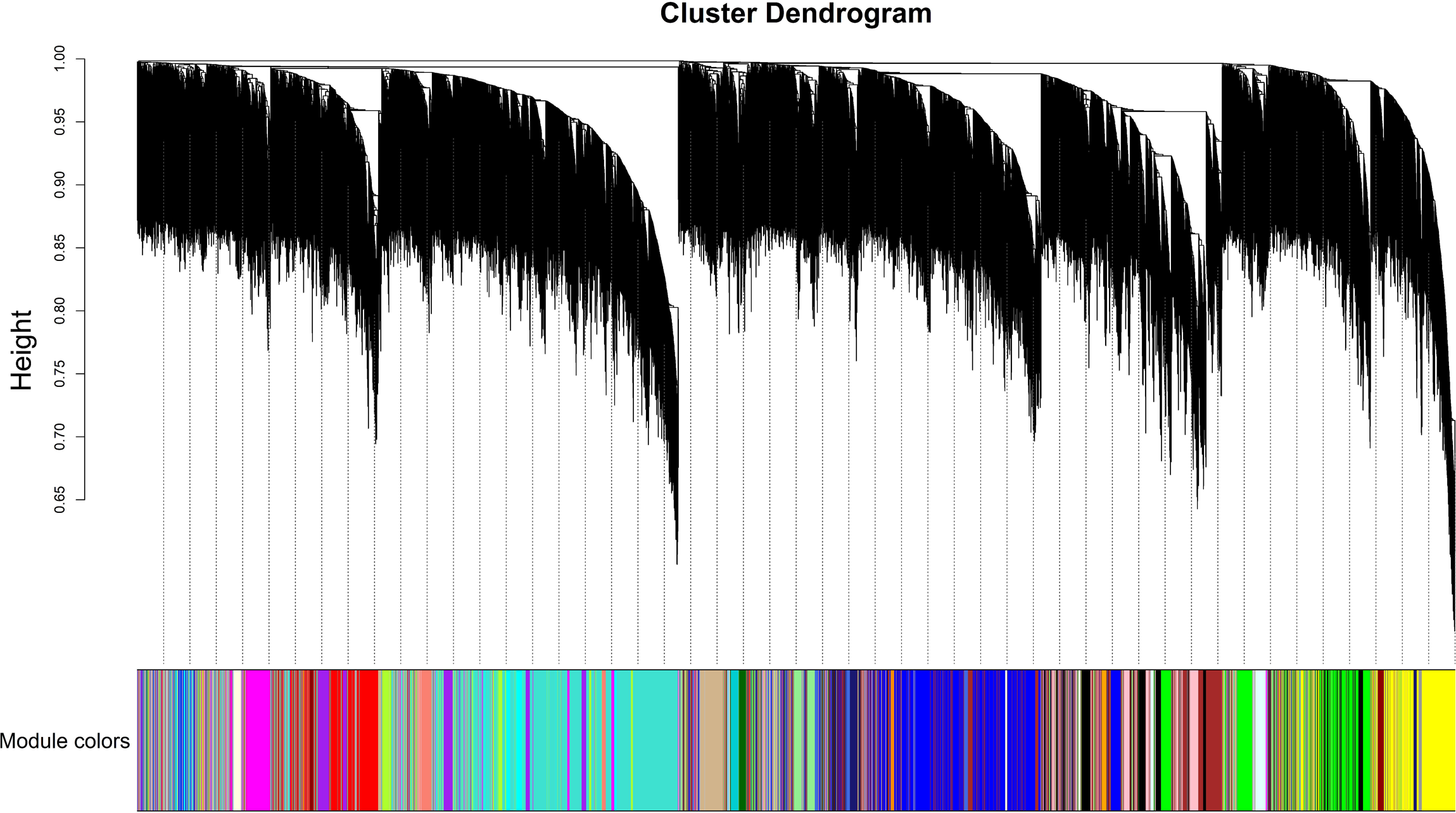
Gene cluster dendrogram based on topological overlap. The colours show the module assignment determined by the Tree cut. Each coloured line represents a color-coded module containing a set of highly connected genes.

### 2.6. Gene Ontology (GO) analysis

GO analysis was performed by following the approach described by Żarski et al. (2021). Briefly, for each transcript, a protein RefSeq accession number was obtained. Next, the RefSeq identifiers were used to align the sequences against human protein in Swiss-Prot with BLASTP. After the alignment, only the best match for each protein was retained, which allowed us to retrieve gene names and UniProt accession numbers for successfully aligned proteins, which were further used to perform GO analysis. GO analysis was performed using the ShinyGO online platform (Ge et al., 2020). First, GO analysis was performed for the genes incorporated in the significant modules obtained from the WGCNA, separately for the positively and negatively correlated modules, and the 10 most enriched biological processes were identified (FDR < 0.05). This allowed us to identify clusters that highlighted the most relevant biological processes. Next, the GO terms found in pre- and post-hatching were compared to identify specific traits unique to each period as well as those shared between them and the results were visualized using a Venn diagram (Bardou et al. 2014). Following the categorization of gene modules based on these two groups of zootechnical traits, we performed a comprehensive GO analysis, resulting in the identification of 100 enriched terms (FDR < 0.05). Subsequently, these terms underwent network analysis through hierarchical clustering. This approach facilitated the identification of distinct clusters, which were further characterized by conducting additional enrichment analysis on the gene list associated with each cluster. Through this process, we discerned the biological processes characteristic of each cluster.

### 2.7. Data analysis and statistics

In addition to the measurable zootechnical characteristics (i.e., mortality, length, weight, cannibalism, SBIE), the specific growth rate (SGR) and the Fulton condition factor (K) were calculated. Fulton’s condition factor (*K*) was calculated based on the obtained measurements according to the formula: K = 100 (W/TL^3^), where W= Weight and TL = total length. In addition, SGR (% day−1) was calculated according to the formula: SGR = 100 ((lnWt–lnW0) Δt^−1^), where W0 = mean initial weight of the fish (g), Wt = mean final weight of the fish (g), and Δt = number of days between measurements. The SGR was calculated considering 3 different conditions/approaches: SGRW_TOT, which refers to the entire rearing trial (0DPH to 27DPH), SGRW_1 relative to the weight of larvae from 0DPH to 17DPH and SGRW_2 for larvae from 17 DPH to 27 DPH. The same calculations were made for the length data of the larvae (referred to as SGRL_TOT, SGRL_1, SGRL_2).

After assessing the normal distribution and homogeneity of variance, one-way, two-way analysis of variance (ANOVA) and Tukey’s post hoc test was conducted; additionally, Kruskal□Wallis’ test and Dunn-Bonferroni post hoc correction were used for analysis data that were not normally distributed. The samples were considered significantly different when *p* < 0.05. The statistical analysis was performed using R studio software ver. 4.2.3 (Team, 2021).

Spearman’s correlation matrix was used in order to evaluate the relationships between the zootechnical traits collected for all the groups. The matrix was constructed in R studio using the “corrplot” package v. 0.92 (Wei et al., 2017).

### 2.8. Key traits for aquaculture (KTA)

The GS values obtained from the WGCNA were used for an additional analysis. The GS is the correlation of gene expression profile with an external trait. It quantifies the biological importance of genes, higher absolute GS values indicate greater significance. GS can be positive or negative. Here, 7 larvae traits relevant for the success of the aquaculture sector were chosen (i.e., mortality, cannibalism, SBIE, SGRL_TOT, SGRW_TOT, K and weight of larvae at the end of the rearing) and the most correlated genes (|cor| > 0.7) highlighted (Supplementary file S3).

Subsequently, the top positively and negatively correlated genes for each traits were validated through real-time quantitative PCR (qPCR). The results were then associated to the specific zootechnical data to assess their correlation coefficient (a value between −1 and +1).

### 2.9. Reverse Transcription and Real-Time qPCR

For this purpose, total RNA was reverse transcribed using a TranScriba kit (A&A Biotechnology, Poland) with oligo(dT)18 primers according to the manufacturer’s instructions. Briefly, 1 µg of total RNA was mixed with 4 µl of 5X reaction buffer, 0.5 µl of RNAse inhibitor, 2 µl of dNTP mix and 4 µl of TranScriba reverse transcriptase. The reaction was conducted for 60 min at 42°C and then completed by heating at 70°C for 5 min.

Real-time qPCR was performed using RT□PCR Mix SYBR (A&A Biotechnology, Poland). For each qPCR (20□µl), 10 ng cDNA template was used along with 10 µl of RT PCR Mix SYBR, 0.5 μM forward and reverse primers (designed with the Primer3Plus online platform (Untergasser et al., 2007) – Supplementary file S4), 0.4 µl of HiRox and sterile water. The reactions were conducted using ViiA7 real-time PCR systems (Applied Biosystems) with the following conditions: incubation at 95°C for 10□min, followed by 40 cycles of denaturation at 95°C for 15□s and annealing and elongation at 60°C for 1□min. After amplification, the efficiency of each primer was calculated using the Real-time PCR Miner program (Zhao and Fernald, 2005). Then, the changes in gene expression were analyzed using the delta delta Ct (2–ΔΔCt) method (Schmittgen and Livak, 2001) as a reference for the geometric mean of four reference genes, namely, acyl-CoA dehydrogenase long chain (*acadl*), thioredoxin 2 (*txn2*), glutathione s-transferase alpha 1 (*gsta1*), and wd repeat domain 83 opposite strand (*wdr83os*), which exhibited the most stable expression level (revealed based on the transcriptomic data obtained) (Żarski et al., 2021).

## 3. Results

All the supplementary figures are gathered in the Supplementary file S5 and are referenced hereinafter as Fig. S1-S14.

### 3.1. Zootechnical performance of larvae

Fertilization rate in all fish larval groups varied between 63% and 97%. Surprisingly, these data didn’t show any significant difference between all the 16 groups (marked as F1 to F16 in Fig.4). Nevertheless, F9 was found to have significantly lower (p < 0.05) embryo developmental rate (after maternal-to-zygotic transition -MZT-) compared to F1 and F6. In addition, two-way ANOVA, which compared the pre- and post-MZT period for each group of larvae, revealed considerable post-MZT mortality in only two groups of larvae (F9 and F10; Fig.4), indicating relatively high consistency between the fertilization rate and post-MZT embryonic survival.

**Fig. 4:**
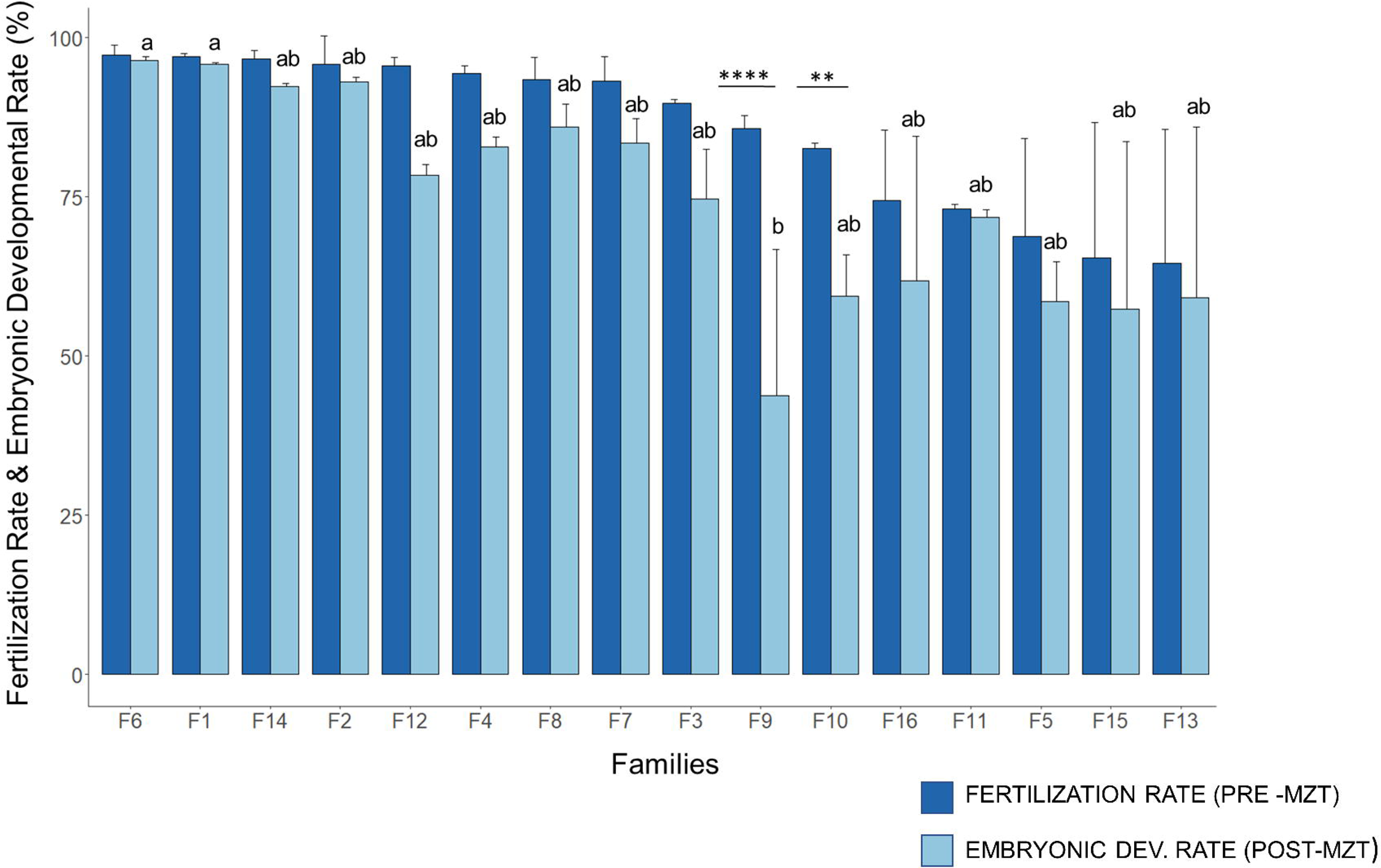
Fertilization rate and embryonic developmental rate of 16 groups of E. perch. The data are arranged in descending order according to the mean values of the fertilization rate between which no significant differences (p>0.05) were recorded. Letters indicate significant differences (p<0.05) between the groups for the embryonic development rate. The asterisks (** – *p <* 0.01, ****– *p <* 0.0001) indicate significant differences within the groups and between fertilization and embryonic development rate.

Among the remaining zootechnical traits, the analysis revealed significant differences (p < 0.05) between groups for mortality, cannibalism (Fig. S1A,B), length and weight of larvae throughout almost the entire rearing period and SGR (for both length and weight) (Fig. S2-S5). However, there was no significant difference for and weight of larvae at mouth opening and first feeding stage and also for the SBIE (Fig. S1C; S3A,B). Also no statistical differences (p > 0.05) were observed for K across different developmental stages (Fig. S6), except for larvae at the end of the experiment.

During the entire rearing trial, the mortality rate of each experimental group was also recorded daily. The cumulative mortality graph (Fig. 5) shows a peak of mortality in mostly all larval groups at oil droplet reduction stage.

**Fig. 5:**
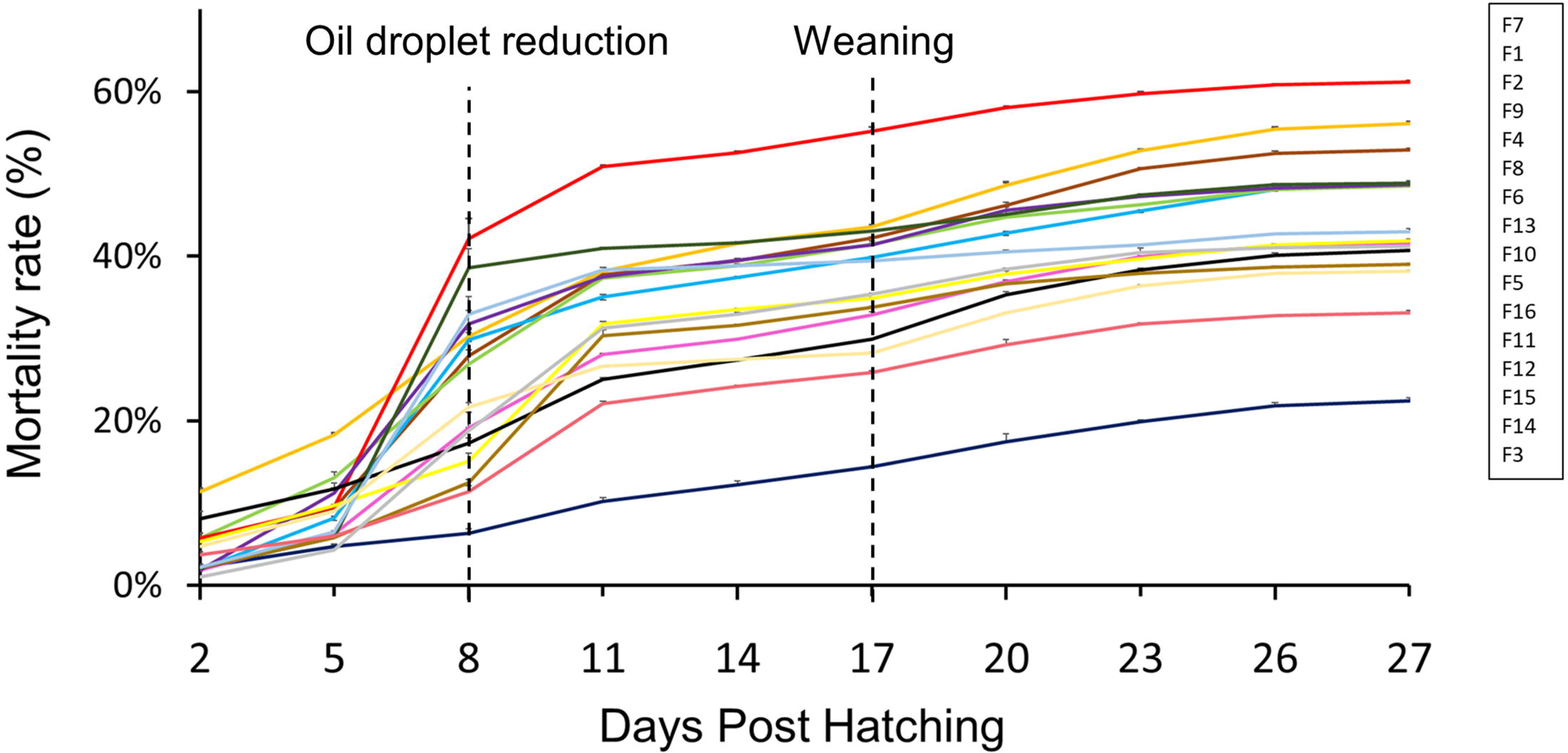
Cumulative mortality (mean ± SD) of 16 groups of E. perch larvae. Different colors stand for different groups. The latter are sorted in descending order of final mortality rate recorded (they are listed in the box at the top right of the graph).

To identify relationships among zootechnical parameters, a Spearman’s correlation (r_s_) matrix was constructed (Fig.6). Interestingly positive correlations between the embryonic developmental rate parameter and fertilization/hatching rates (respectively r_s_ = 0.89 and r_s_ = 0.7) are shown. Noteworthy, robust correlations (r_s_ ≥ 0.6 or r_s_ ≥ −0.6) are observed for the growth-related traits. For instance, length of the larvae at mouth opening correlates positively with length at the first feeding (r_s_ = 0.67), weight at first feeding (r_s_ = 0.66) and at oil droplet reduction (r_s_ = 0.63). Specific growth rate of weight for the entire larviculture period (SGRW_TOT) is negatively correlated with larval weight at mouth opening stage (r_s_ = - 0.61), while it is positively related to weight and length of larvae at the end of the experiment (respectively, r_s_ = 0.86, r_s_ = 0.73). Alternatively, the negative relationship between *K* at weaning (Fulton_W) and the specific growth rate for weight data from weaning to the end of the larval period (SGRW_2) (r_s_ = - 0.65). Also, the matrix shows negative correlations between *K* at mouth opening (Fulton_MO) and fertilization and embryonic developmental rate (r_s_ = −0.65, r_s_ = −0.63) (Fig. 5). This can be interpreted as the larvae obtained from lower quality eggs appear to be much more robust, resulting in a better *K*.

**Fig. 6:**
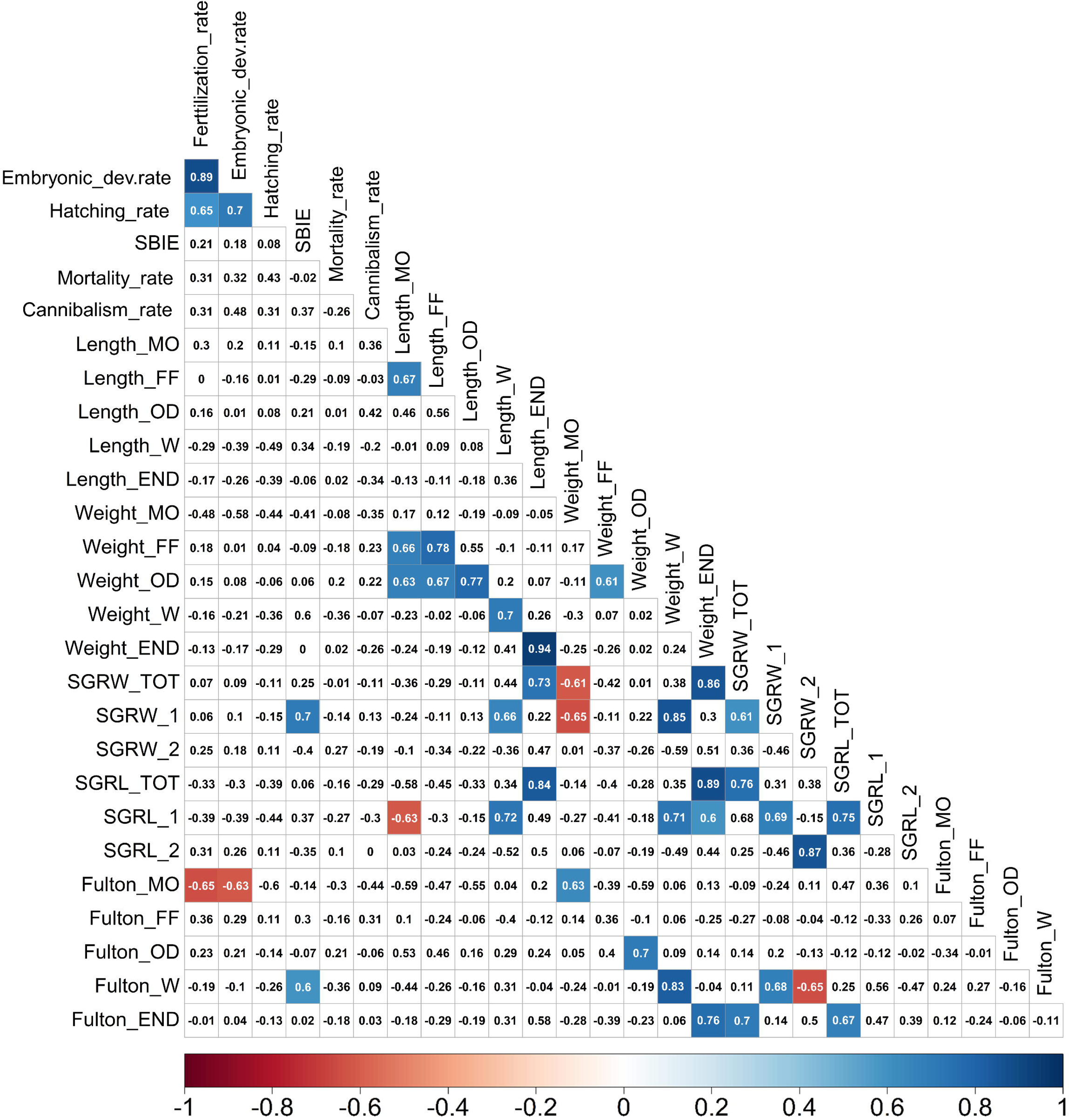
Spearman’s correlation matrix of all the zootechnical traits and for all the groups together. Correlation coefficients (r_s_) are shown in the squares, with significant correlations (r_s_ ≥ 0.6, p < 0.05) indicated by coloured boxes. Red colours show significant negative correlations, blue shows significant positive correlation, and white shows insignificant correlations. SBIE: swim bladder inflation effectiveness, MO: mouth opening, FF: first feeding, OD: oil droplet reduction, W: weaning, END: end of the experiment, SGRW: specific growth rate for weight, SGRL: specific growth rate for length data, SGR_TOT: specific growth rate for the entire larviculture period, SGR_1: specific growth rate from hatching until weaning stage, SGR_2: specific growth rate from weaning stage until the end of the experiment, Fulton: Fulton’s condition factor.

### 3.2. Transcriptomic data and WGCNA analysis

After RNA-sequencing, 30,744 genes were initially identified. Following filtering procedures (explained in paragraph 2.5 in Material and Methods), 19,656 genes were analysed utilizing WGCNA. Initially, a gene cluster dendrogram was constructed, yielding 28 distinct gene modules (Fig. 3). Next, correlation analysis between these modules and zootechnical traits are shown in the module-traits heatmap (Fig. 7). Notably a total of 13 modules with p < 0.05 and correlation coefficient |cor| ≥ 0.6 were prioritized for further scrutiny. Specifically, 7 modules exhibited notably robust correlations (r ≥ 0.6) with embryonic developmental rate and hatching rate, indicating their association with pre-hatching traits. Conversely, 9 modules displayed significant correlations with post-hatching traits, specifically the weight of larvae at mouth opening, length at first feeding, K at mouth opening and oil droplet reduction stage. Notably, modules black, pink, and turquoise demonstrated shared correlations with both pre-hatching and post-hatching traits, suggesting their importance across different developmental stages.

**Fig. 7:**
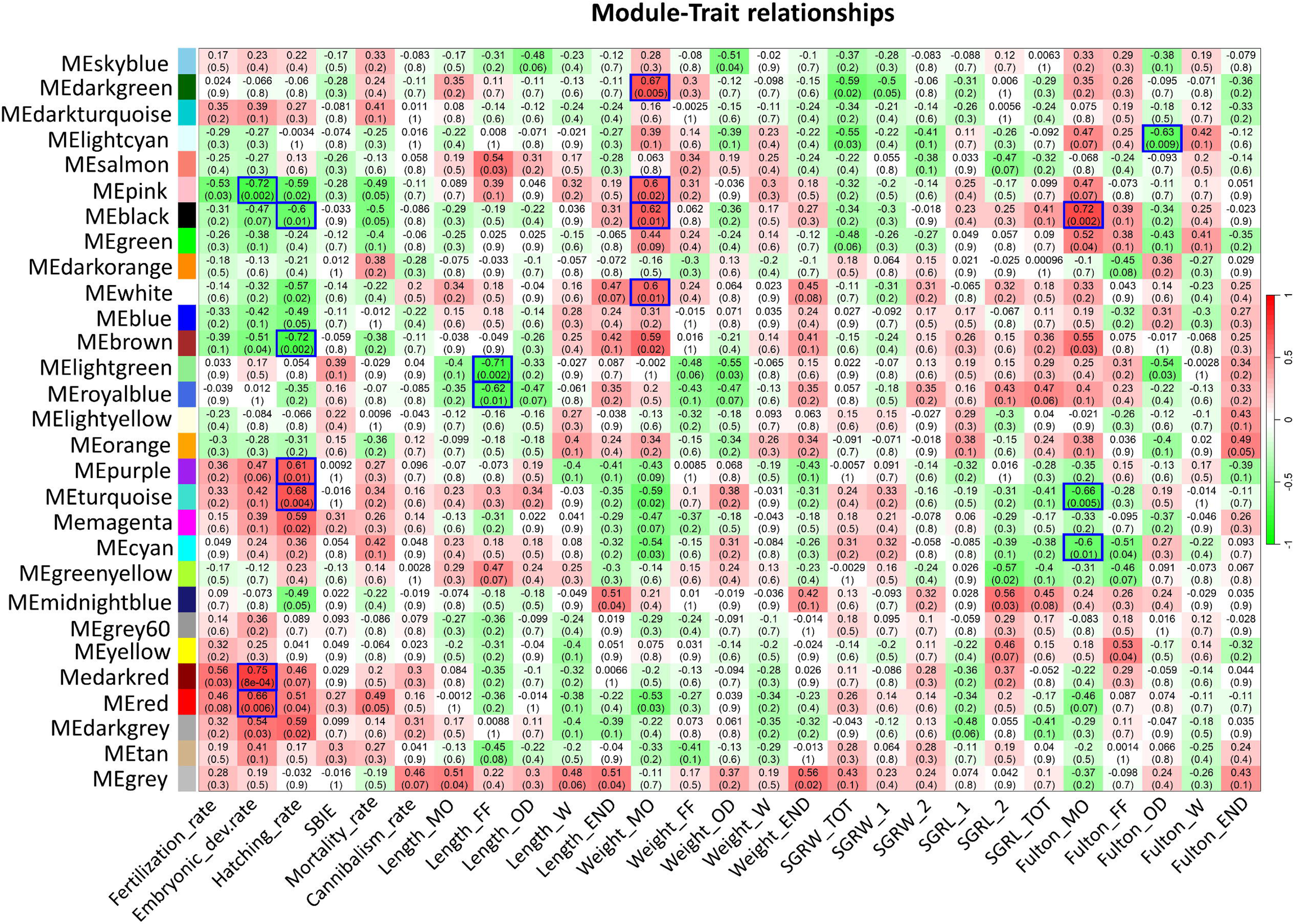
Module-traits relationship. The module eigengene (ME) is shown in each row. Genes not assigned to any of the other modules are included in the grey module. The columns represent the zootechnical traits. The modules with high correlation values and *p <* 0.05 were identified as significant trait-related modules. The colors indicate the positive (red) and negative (green) correlations between gene modules and traits. SBIE: swim bladder inflation effectiveness, MO: mouth opening, FF: first feeding, OD: oil droplet reduction, W: weaning, END: end of the experiment, SGRW: specific growth rate for weight, SGRL: specific growth rate for length data, SGR_TOT: specific growth rate for the entire larviculture period, SGR_1: specific growth rate from hatching until weaning stage, SGR_2: specific growth rate from weaning stage until the end of the experiment, Fulton: Fulton’s condition factor.

Subsequent Gene Ontology (GO) Enrichment analysis was performed for those 13 modules. This analysis revealed enriched biological processes associated with specific traits. Regarding the embryonic developmental rate, negatively correlated modules were associated with terms linked to ribosome biogenesis, RNA processing, and modification (Fig. S7A). Conversely, positively correlated modules were enriched in terms related to morphogenesis, circulatory system development, and response to endogenous stimuli (Fig. S7B). For hatching rate, GO analysis indicated a positive correlation with processes primarily involved in neurogenesis, while negatively correlated processes included those related to protein transport and metabolic processes such as organic acid and carboxylic acid metabolism (Fig. S8A,B). Modules significantly correlated with larval weight at mouth opening were exclusively positively associated, with GO analysis implicating mechanisms related to RNA processing and ribosome biogenesis (Fig. S9). In contrast, length at first feeding showed negative correlations with two modules, revealing processes associated with neurotransmitter transport, transmembrane transport, ion transport, and chemical synaptic transport (Fig. S10). The analysis of negative modules for Fulton’s condition coefficient at mouth opening were associated with genes related to neurogenesis, neuron generation, and differentiation (Fig. S11A), while positive modules for Fulton’s condition coefficient at mouth opening indicated involvement in translation, ribosome biogenesis, and metabolic processes (Fig. S11B). Furthermore, significant modules were identified for K at mouth opening and the oil droplet reduction phase. The latter negatively correlated with processes involved in the mitotic cell cycle, chromosome segregation, and nuclear division (Fig. S12).

In summary, it is crucial to note that the transcriptomic profile observed at the larvae’s mouth opening stage is shaped by genes that correlate with traits seen before and after hatching, encompassing a variety of functions. Among the enriched biological processes, those related to regulation of transcription processes, cell differentiation and signal transduction are linked to events occurring before hatching. In contrast, processes associated with mitotic cell cycle are specific of traits manifesting after hatching (Fig. S13B). Intriguingly, there are common biological processes identified for both pre- and post-hatching traits, particularly those related to neurodevelopment (Fig. S13C). This suggests that neurodevelopment plays a key role in driving both embryonic and larval development stages.

### 3.3. Key traits for aquaculture (KTA)

WGCNA employs significance calculations to pinpoint genes intricately associated with targeted traits. These findings facilitate the identification of most correlated genes for traits such as mortality, cannibalism, SBIE, SGR for total weight and length, K and weight of larvae at the end of the experimental period. The results unveil a comprehensive table showcasing genes with high correlations to each examined parameter (Supplementary file S3), offering valuable insights into the molecular underpinnings of these traits. The qPCR validation confirmed the association of specific genes, such as *selenoprotein O (selenoo), tripartite motif-containing protein 16 (trim16), solute carrier family 15 member 1 (slc15a1), clock-interacting pacemaker (cipc)*, with the traits under investigation, as outlined in Table 1 (see also Fig. S14). These genes are constituting candidate markers which could serve to predict the traits they are significantly correlated with.

**Table 1:** Most correlated genes. (both positive and negative) selected for each selected commercially relevant traits, further validated with qPCR. The resulting table includes the gene significance (GS) values obtained from weighted gene co-expression network analysis (WGCNA), correlation values from qPCR. SBIE: swim bladder inflation effectiveness, SGRW_TOT: specific growth rate for weight for the entire larviculture period, SGRL_TOT: specific growth rate for length for the entire larviculture period, Fulton_END: Fulton’s condition factor at the end of the experiment.

|  | CANNIBALISM |  | MORTALITY |  | SBIE |  | WEIGHT END |  | SGRW_TOT |  | SGRL_TOT |  | FULTON_END |  |
| --- | --- | --- | --- | --- | --- | --- | --- | --- | --- | --- | --- | --- | --- | --- |
|  | *selenoo* | *nudt12* | *mkx* | *crcp* | *si:dkey-117m1.4* | *pycard* | *aoc1* | *cipc* | *prdm1* | *txnl4a* | *atp8a2* | *slc15a1* | *sec14l2* | *trim16* |
| GS | 0.82 | -0.85 | 0.77 | -0.81 | 0.79 | -0.78 | 0.81 | -0.82 | 0.84 | -0.84 | 0.87 | -0.80 | 0.81 | -0.80 |
| qPCR | 0.78 | -0.31 | -0.02 | -0.18 | -0.32 | -0.33 | -0.02 | -0.70 | 0.01 | -0.24 | 0.13 | -0.72 | 0.04 | -0.55 |

## 4. Discussion

Studies conducted on the early developmental stages of larvae can provide valuable insights into the factors that influence, and consequently predict their future performance. Despite advancements, our comprehension of the mechanisms responsible for successful development and survival remains limited. By combining traditional zootechnical assessments with transcriptomic analyses, our research yielded significant insights into larval biology paving the way for understanding intricate growth changes and molecular mechanisms that contribute to the performance of fish larvae. Examining 16 distinct larval groups, originating from diverse parental pairs, facilitated an in-depth exploration of larval transcriptomes. The data presented draws our attention to the fact that the molecular profile of freshly hatched larvae (at mouth-opening stage) is highly indicative of pre-hatching events. This emphasize crucial role of parental contribution in shaping the larval transcriptomic landscape. However, our findings shed also light on the importance of the larvae’s transcriptomic profile in determining their future performance, being a solid foundation for further investigations.

### 4.1. Zootechnical traits

Overall, the observed growth patterns (in terms of weight and length) align with previous studies on Eurasian perch (Kupren et al., 2019; Palińska-Żarska et al., 2020). Additionally, there were no major differences observed in cannibalism intensity and SBIE when compared to previous experiments that exposed the larvae to similar experimental conditions (Kupren et al., 2019; Palińska-Żarska et al., 2020). In addition, larval mortality increased especially at the oil droplet reduction phase, reflecting the challenges encountered by the larvae when they switch entirely to exogenous feeding (Kestemont et al., 2003; Król et al., 2019; Palińska-Żarska et al., 2020). This phenomenon could be linked to the non-feeding behaviour, commonly observed in fish larval species (Yúfera and Darias, 2007).

Upon closer observation of the data obtained, although all groups were reared under the same controlled conditions, considerable variation between experimental groups in terms of zootechnical traits was observed, which reflected different performances. Growth-related differences likely influenced cannibalistic tendencies and overall fish development. Such growth heterogeneity has been commonly detected in many other fish species and has been always considered as a relevant determinant and predictor of fish survival, since it can lead to aggressiveness and mortality (Baras and Dalmeida, 2001; Carvalho et al., 2018; Kestemont et al., 2003).

The correlation analysis between various zootechnical traits provides a comprehensive overview of the interrelationships among qualitative traits, elucidating the overall growth trajectory of the offspring. Notably, a positive correlation between embryonic developmental rate and hatching rate suggests that embryos which passed the MZT mostly hatched successfully, supporting the significance of embryonic developmental rate as a reliable indicator of egg developmental competence (Bobe, 2015). Additionally, the analysis illustrates a positive correlation between the length and weight of fry at each sampling time point, indicating that larvae tend to become more robust as their length increases. Conversely, negative correlations, such as those observed between the weight of larvae at mouth opening and SGR from hatching until weaning stage and at the experiment’s end, may signify a sort of compensatory mechanism in growth between early and later developmental stages. This phenomenon is commonly observed and has been documented for various fish species (Ali et al., 2003).

The observed correlations between embryonic developmental rates, fertilization and hatching rates, and between growth-related traits, provide insights into larval development dynamics and confirms that these parameters as valuable predictors of future fish performance and overall fitness (Brooks et al., 1997; Koumoundouros et al., 2017). Also, the variability observed in zootechnical traits despite identical rearing conditions implies the involvement of inherited parental factors shaping larvae features. This underscores the need to investigate the molecular background of larvae to elucidate this phenomenon.

### 4.2. Transcriptome data analysis - between past and future

WGCNA identifies gene modules linked to larval traits, aiming to pinpoint specific zootechnical indicators and their dependency with the larval transcriptome as well as to reveal underlying molecular mechanisms crucial in the early phases of fish growth. The results underscored significant correlation between gene networks and pre-hatching parameters, as embryonic developmental and hatching rates, as well as post-hatching traits, i.e. weight at mouth opening, length at first feeding, K at mouth opening and at oil droplet reduction stage. Overall, the GO enrichment analysis of the genes identified within modules exhibit diverse functional processes, including cell cycle, RNA processing, ribosome biogenesis, protein trafficking, apoptosis, circulatory system regulation, and neurogenesis. The consistency of these findings across different fish species (Mazurais et al., 2011; Bougas et al., 2013) further emphasizes the significance of these processes in shaping larval performance and underscores also its complexity in fish.

Embryonic developmental rate and hatching rate, were negatively correlated to genes involved in RNA processing, translation, ribosome biogenesis and protein transports. These processes collectively govern gene expression, cell signalling, and tissue differentiation, profoundly impacting fish larval growth and survival. Particularly, ribosomes produced abundantly during oogenesis and presumably deposited in the eggs, play a crucial role in synthesizing proteins vital for various developmental processes (Leesch et al., 2023; Qi et al., 2016; Shen et al., 2017). These maternally provided ribosome are paramount during embryogenesis specifically until Zygotic Genome Activation (ZGA), which marks the transition of developmental control from maternal to zygotic factors (Leesch et al., 2023). Since the pre-hatching traits were negative linked to this biological processes, our speculation is that post-ZGA, embryos likely redirect their energy towards processes -positively correlated with pre-hatching traits – like neurogenesis, sensory organ development, and morphogenesis of tubes and blood vessels, which are equally necessary for their future developmental success. These results indicate that embryos may prioritize these processes closer to hatching. The embryos, as well as post-hatching larvae, are constantly subjected to morphological and physiological modifications. Among others, the maturation of the nervous system is one of the most important event. This process is crucial for enhancing sensory perception, motor coordination, and cognitive functions in larvae (Nelson and Granato, 2022). Previous studies discussed the nervous systems’ development could potentially be pre-programmed by the molecular content inherited maternally (Żarski et al., 2021, 2020b). This, along with environmental factors may influence the trajectory of nervous system development and impact future behaviour and adaptive responses (Colson et al., 2019). In summary, right after fertilization more general but crucial functions are prioritized (i.e. ribosome biogenesis), while after ZGA processes essential for successful accomplishment of embryonic development take precedence. These subsequent events will most likely define the success of the hatching.

Molecular profile of the larvae is determined by the interplay between environmental factors, genetic background and so called non-genetic inheritance mechanisms. The latter encompass, among others, mRNAs which play a pivotal role as modulators of gene expression during the early development and consequently influencing the phenotype of the progeny (Adrian-Kalchhauser et al., 2020). It has been hypothesized that such cascade-like transmission of information from parent to progeny affects the performance of larvae and juveniles (Adrian-Kalchhauser et al., 2018; Colson et al., 2019). In this context, results of our study highlights the importance of parental contribution in shaping the transcriptomic profile of the larvae (Harvey et al., 2013), reflecting their past. Analogically, the significant correlations with the post hatching traits are predictive of the larvae’s future. Notably, those traits fall within the initial growth phases until yolk sac absorption. Up to this stage, larvae rely primarily on the nutrients stored within the yolk sac, which they inherited from the female (Callet et al., 2022). These nutrients play a crucial role in sustaining the larvae’s early development by determining energetic reservoirs and indirectly controlling their growth and performance (Bachan et al., 2012; Migaud et al., 2013). The GO enrichment analysis of gene modules linked to post-hatching traits aligns with molecular processes identified in pre-hatching traits. However, contrarily to pre-hatching traits, genes associated with ribosome biogenesis exhibit a positive correlation with post-hatching traits, while those linked to neurogenesis show a negative correlation. This inversion may suggest a shift in biological process prioritization as larvae transition, depending on their developmental stage (Mathavan et al., 2005). For example, after hatching larvae may prioritize protein production to overcome crucial metamorphosis events (e.g. onset of exogenous feeding, swim bladder inflation, etc). We can hypothesize that bigger larvae at mouth opening invest more in protein translation, potentially leading to enhanced physiological development and growth rates. Also, upon first feeding larvae that exhibit larger sizes, may have already better-developed senses and a more advanced nervous system, which can explain the inverse correlation with neurogenesis pathways at this stage.

Post-hatching traits exhibit a specific association with processes linked to the cell cycle and mitosis. As larvae undergo metamorphosis and confront various challenges related to the interaction with the external environment, significant restructuring of organs and tissues occurs, with the cell cycle serving as a central mechanism (González-Quirós et al., 2007). This transition likely initiates a cascade of molecular events, indicating a profound shift in cellular activities towards enhancing the growth and development of various organs to support their functions. For instance, after the oil droplet reduction stage, larvae become entirely dependent on external feed for energy, necessitating the digestive system to efficiently process and utilize food resources. Rapid cell division and proliferation may play critical roles in optimizing system functions, enabling larvae to better adapt and respond to environmental cues.

Overall, the contrasting correlations between ribosome biogenesis and neurogenesis well reflect the dynamic nature of larval development. As larvae progress through various stages, their biological priorities shift, leading to fluctuations in gene expression patterns and molecular processes. Understanding these intricate relationships between molecular processes and larval development provides valuable insights into the adaptive strategies of organisms and the mechanisms underlying developmental plasticity. Nonetheless, the environmental factors still play a significant role in shaping these developmental trajectories. Changes in temperature, light exposure, nutrient availability, and other environmental cues can modulate gene expression and influence the balance between different biological processes (Mazurais et al., 2011; Urho, 2002). Our results indicates that transcriptomic signature of larvae at the mouth opening stage offers significant insights into parental contributions and their influence on embryogenesis. However, its ability to predict future larval performance appears to be somewhat limited. It should be emphasized, that future larval fate is determined by the interaction of their molecular cargo with the external factors playing a major role right after hatching. It’s essential to acknowledge that in our ‘common garden’ experiment all the environmental factors were controlled and equal for all groups of larvae. This may limit the detection of certain traits typically observed in wild conditions, such as exposure to pathogens or stressors. For instance, immune system traits may not be evident in our current setup, but exposing organisms to bacterial or temperature challenges could reveal the transcriptome’s predictive capacity for stress or immune system functioning, which is important in fish adaptability (Elabd et al., 2017; Kammer et al., 2011; Palińska-Żarska et al., 2021). Therefore, future research should consider subjecting organisms to specific challenges to uncover additional predictive traits.

### 4.3. Key traits for aquaculture (KTA) - molecular signatures

Aquaculture relies on important traits like cannibalism, survival rate, swim bladder inflation, weight, and specific growth rate for successful management and production (Toomey et al., 2021). The transcriptome of newly hatched larvae may offer predictive insights into future development, supporting aquaculture research and its production. The WGCNA has facilitated the identification of genes strongly associated with these commercially relevant traits, potentially serving as specific gene markers.

The validation with qPCR highlighted 4 genes: *selenoo, trim16, slc15a1, cipc*; linked respectively to cannibalism, SGR for length, K and weight of larvae at the end of the experiment. Notably, *cipc* appears pivotal in regulating various physiological processes in fish, influencing behaviour, metabolism, and clock genes seem to be implicated in thermal resistance (Hung et al., 2016); while *slc15a1* affects nutrient absorption, transport and fish growth(Romano et al., 2014; Vacca et al., 2019). *Trim16*, a member of the teleost-specific *fintrim* family (van der Aa et al., 2009), is involved in regulation of innate immunity, but it seems also involved in cell proliferation, differentiation, and metabolism (Cho et al., 2022). While its specific functions in fish larvae, particularly regarding traits like Fulton’s condition factor, may require further investigation, its involvement in cellular processes suggests potential roles in larval development and physiology. Also, selenoproteins, including *selenoo*, contribute to cellular antioxidant defence mechanisms (Han et al., 2014; Sumana et al., 2023), and indirectly, they can play a role in fish growth.

These genes collectively play crucial roles in fish larval development and physiology, impacting growth, behaviour, immunity, and metabolic processes. However, out of the 14 chosen genes, only 4 were confirmed through qPCR validation. This suggests that our approach (i.e. finding gene markers based on correlation of their expression with traits) may not be entirely reliable for detecting gene markers associated with zootechnical traits. Future studies should consider employing more pertinent approaches, such as conducting specific experiments tailored to assess each trait individually stimulating higher variability in the expression. This would provide a clearer understanding of the mechanisms and interactions underlying larval traits and overall fish performance.

## 5. Conclusion

Considering all the above mentioned, larval transcriptomic profile represents a bridge between the past, stemming from the transcripts provided by parents within gametes and continuing through embryonic development, and the future, serving as a footprint for forthcoming instructions for the larval period and potentially beyond. Taking all of this into account, by analysing transcriptome of freshly hatched larvae we gather information on parentally derived molecular cargo but also on larvae adaptability, paving the way to comprehend the intricate developmental trajectories that lead to adulthood.

## Acknowledgments

The authors would like to acknowledge Dr Jarosław Król for his valuable help with logistics. Additionally, Prof. Stefan Dobosz and Rafał Rożyński for providing facilities to maintain the fish overwinter.

## Competing interests

The authors declare that they have no competing interests.

## Authors’ Contribution

**Rossella Debernardis**: Conceptualization and conduction of the experiment, samples collection, data analysis, data curation, writing - original draft. **Katarzyna Pali**ń**ska-**Ż**arska** Conceptualization and conduction of experiment, samples collection, Writing - review & editing, Supervision. **Sylwia Judycka:** Conceptualization and conduction of the experiment, Investigation. **Abhipsa Panda:** Investigation. **Sylwia Jarmołowicz:** Investigation. **Jan P. Jastrz**ę**bski**: data curation. **Tainá Rocha de Almeida**: data curation**. Maciej Bła**ż**ejewski:** Resources. **Piotr Hliwa:** Resources**. Sławomir Krejszeff:** Resources. **Daniel** Ż**arski**: Conceptualization, Funding acquisition, Project administration, Supervision, Writing – review & editing. All authors read and approved the final manuscript.

## Funding

This research was a part of the project “Transcriptomic and zootechnical exploration of parental contribution to progeny quality in Eurasian perch, *Perca fluviatilis*” funded by the National Science Centre of Poland (SONATA BIS project, number UMO-2020/38/E/NZ9/00394).

## Availability of data and materials

Raw data from the analysis of different groups of freshly hatched larvae can be accessed via the NCBI BioProject database under the PRJNA1032718 accession number (restricted availability prior publication).

## Notes

### Competing Interest Statement

The authors have declared no competing interest.

### Summary of Updates

This version of the MS has been revised following critical reviews received by Authors from a scientific journal, which were related with the interpretation of the data. This has led us to revision of the paper into current form, which is now also meeting the criticism received during the peer-review process. So, in effect we have modified the concept of the study, changed the title and substantially revised the paper.

## References

Adrian-Kalchhauser, I., Sultan, S.E., Shama, L.N.S., Spence-Jones, H., Tiso, S., Keller Valsecchi, C.I., Weissing, F.J., 2020. Understanding “Non-genetic” Inheritance: Insights from Molecular-Evolutionary Crosstalk. Trends Ecol Evol. 10.1016/j.tree.2020.08.011

Adrian-Kalchhauser, I., Walser, J.C., Schwaiger, M., Burkhardt-Holm, P., 2018. RNA sequencing of early round goby embryos reveals that maternal experiences can shape the maternal RNA contribution in a wild vertebrate. BMC Evol Biol 18. 10.1186/s12862-018-1132-2

Ali, M., Nicieza, A., Wootton, R.J., 2003. Compensatory growth in fishes: A response to growth depression. Fish and Fisheries 4. 10.1046/j.1467-2979.2003.00120.x

Bachan, M.M., Fleming, I.A., Trippel, E.A., 2012. Maternal allocation of lipid classes and fatty acids with seasonal egg production in Atlantic cod (Gadus morhua) of wild origin. Mar Biol 159. 10.1007/s00227-012-2014-6

Baras, É., Dalmeida, A.F., 2001. Size heterogeneity prevails over kinship in shaping cannibalism among larvae of sharptooth catfish Clarias gariepinus. Aquat Living Resour 14. 10.1016/S0990-7440(01)01118-4

Bobe, J., 2015. Egg quality in fish: Present and future challenges. Animal Frontiers 5. 10.2527/af.2015-0010

Boglione, C., Gavaia, P., Koumoundouros, G., Gisbert, E., Moren, M., Fontagné, S., Witten, P.E., 2013a. Skeletal anomalies in reared European fish larvae and juveniles. Part 1: Normal and anomalous skeletogenic processes. Rev Aquac. 10.1111/raq.12015

Boglione, C., Gisbert, E., Gavaia, P., Witten, P.E., Moren, M., Fontagné, S., Koumoundouros, G., 2013b. Skeletal anomalies in reared European fish larvae and juveniles. Part 2: Main typologies, occurrences and causative factors. Rev Aquac 5. 10.1111/raq.12016

Bolger, A.M., Lohse, M., Usadel, B., 2014. Trimmomatic: A flexible trimmer for Illumina sequence data. Bioinformatics 30. 10.1093/bioinformatics/btu170

Bougas, B., Audet, C., Bernatchez, L., 2013. The influence of parental effects on transcriptomic landscape during early development in brook charr (Salvelinus fontinalis, Mitchill). Heredity (Edinb) 110. 10.1038/hdy.2012.113

Brooks, S., Tyler, C.R., Sumpter, J.P., 1997. Egg quality in fish: What makes a good egg? Rev Fish Biol Fish 7. 10.1023/A:1018400130692

Buchfink, B., Reuter, K., Drost, H.G., 2021. Sensitive protein alignments at tree-of-life scale using DIAMOND. Nat Methods 18. 10.1038/s41592-021-01101-x

Callet, T., Cardona, E., Turonnet, N., Maunas, P., Larroquet, L., Surget, A., Corraze, G., Panserat, S., Marandel, L., 2022. Alteration of eggs biochemical composition and progeny survival by maternal high carbohydrate nutrition in a teleost fish. Sci Rep 12. 10.1038/s41598-022-21185-5

Cantalapiedra, C.P., HernLJandez-Plaza, A., Letunic, I., Bork, P., Huerta-Cepas, J., 2021. eggNOG-mapper v2: Functional Annotation, Orthology Assignments, and Domain Prediction at the Metagenomic Scale. Mol Biol Evol 38. 10.1093/molbev/msab293

Carvalho, T.B., de Souza, E.C.M., Pinheiro-Da-Silva, J., Villacorta-Correa, M.A., 2018. Effect of body size heterogeneity on the aggressive behavior of larvae of matrinxã, Brycon amazonicus (Characiformes, Bryconidae). Acta Amazon 48. 10.1590/1809-4392201800541

Chandhini, S., Rejish Kumar, V.J., 2019. Transcriptomics in aquaculture: current status and applications. Rev Aquac. 10.1111/raq.12298

Cho, J.Y., Kim, J., Kim, J.W., Lee, D., Kim, D.G., Kim, Y.S., Lee, J.H., Nam, B.H., Kim, Y.O., Kong, H.J., 2022. Characterization of TRIM16, a member of the fish-specific finTRIM family, in olive flounder Paralichthys olivaceus. Fish Shellfish Immunol 127. 10.1016/j.fsi.2022.07.003

Colson, V., Cousture, M., Damasceno, D., Valotaire, C., Nguyen, T., Le Cam, A., Bobe, J., 2019. Maternal temperature exposure impairs emotional and cognitive responses and triggers dysregulation of neurodevelopment genes in fish. PeerJ 2019. 10.7717/peerj.6338

Dobin, A., Davis, C.A., Schlesinger, F., Drenkow, J., Zaleski, C., Jha, S., Batut, P., Chaisson, M., Gingeras, T.R., 2013. STAR: Ultrafast universal RNA-seq aligner. Bioinformatics 29. 10.1093/bioinformatics/bts635

Elabd, H., Wang, H.P., Shaheen, A., Yao, H., Abbass, A., 2017. Anti-oxidative effects of some dietary supplements on Yellow perch (Perca flavescens) exposed to different physical stressors. Aquac Rep 8. 10.1016/j.aqrep.2017.09.002

Ferraresso, S., Bonaldo, A., Parma, L., Cinotti, S., Massi, P., Bargelloni, L., Gatta, P.P., 2013. Exploring the larval transcriptome of the common sole (Solea solea L.). BMC Genomics 14. 10.1186/1471-2164-14-315

Fontaine, P., Teletchea, F., 2019. Domestication of the Eurasian Perch (Perca fluviatilis), in: Animal Domestication. 10.5772/intechopen.85132

Ge, S.X., Jung, D., Jung, D., Yao, R., 2020. ShinyGO: A graphical gene-set enrichment tool for animals and plants. Bioinformatics 36. 10.1093/bioinformatics/btz931

González-Quirós, R., Munuera, I., Folkvord, A., 2007. Cell cycle analysis of brain cells as a growth index in larval cod at different feeding conditions and temperatures. Sci Mar 71. 10.3989/scimar.2007.71n3485

Güralp, H., Pocherniaieva, K., Blecha, M., Policar, T., Pšenička, M., Saito, T., 2016. Early embryonic development in pikeperch (Sander lucioperca) related to micromanipulation. Czech Journal of Animal Science 61. 10.17221/35/2015-CJAS

Han, S.J., Lee, B.C., Yim, S.H., Gladyshev, V.N., Lee, S.R., 2014. Characterization of mammalian selenoprotein O: A redox-active mitochondrial protein. PLoS One 9. 10.1371/journal.pone.0095518

Harvey, S.A., Sealy, I., Kettleborough, R., Fenyes, F., White, R., Stemple, D., Smith, J.C., 2013. Identification of the zebrafish maternal and paternal transcriptomes. Development (Cambridge) 140. 10.1242/dev.095091

Huerta-Cepas, J., Szklarczyk, D., Heller, D., Hernández-Plaza, A., Forslund, S.K., Cook, H., Mende, D.R., Letunic, I., Rattei, T., Jensen, L.J., Von Mering, C., Bork, P., 2019. EggNOG 5.0: A hierarchical, functionally and phylogenetically annotated orthology resource based on 5090 organisms and 2502 viruses. Nucleic Acids Res 47. 10.1093/nar/gky1085

Hung, I.C., Hsiao, Y.C., Sun, H.S., Chen, T.M., Lee, S.J., 2016. MicroRNAs regulate gene plasticity during cold shock in zebrafish larvae. BMC Genomics 17. 10.1186/s12864-016-3239-4

Jastrzebski, J.P., Pascarella, S., Lipka, A., Dorocki, S., 2023. IncRna: The R Package for Optimizing lncRNA Identification Processes. Journal of Computational Biology 30, 1322–1326.

Judycka, S., Żarski, D., Dietrich, M.A., Karol, H., Hliwa, P., Błażejewski, M., Ciereszko, A., 2022. Toward commercialization: Improvement of a semen cryopreservation protocol for European perch enables its implementation for commercial-scale fertilization. Aquaculture 549. 10.1016/j.aquaculture.2021.737790

Judycka, S., Żarski, D., Dietrich, M.A., Palińska-Żarska, K., Karol, H., Ciereszko, A., 2019. Standardized cryopreservation protocol of European perch (Perca fluviatilis) semen allows to obtain high fertilization rates with the use of frozen/thawed semen. Aquaculture 498. 10.1016/j.aquaculture.2018.08.059

Kammer, A.R., Orczewska, J.I., O’Brien, K.M., 2011. Oxidative stress is transient and tissue specific during cold acclimation of threespine stickleback. Journal of Experimental Biology 214. 10.1242/jeb.053207

Kestemont, P., Jourdan, S., Houbart, M., Mélard, C., Paspatis, M., Fontaine, P., Cuvier, A., Kentouri, M., Baras, E., 2003. Size heterogeneity, cannibalism and competition in cultured predatory fish larvae: Biotic and abiotic influences, in: Aquaculture. 10.1016/S0044-8486(03)00513-1

Koumoundouros, G., 2010. Morpho-Anatomical Abnormalities in Mediterranean Marine Aquaculture. Recent Advances in Aquaculture Research.

Koumoundouros, G., Gisbert Enric, Fernandez, I., Cabrita, E., Galindo-Villegas, J., Conceição, L.E.C., 2017. Quality descriptors and predictors in farmed marine fish larvae and juveniles. John Wiley & Sons, Incorporated, ProQuest Ebook Central: Oxford, UK, 2017.

Król, J., Długoński, A., Błażejewski, M., Hliwa, P., 2019. Effect of size sorting on growth, cannibalism, and survival in Eurasian perch Perca fluviatilis L. post-larvae. Aquaculture International 27. 10.1007/s10499-018-00337-3

Kupren, K., Palińska-Żarska, K., Krejszeff, S., Żarski, D., 2019. Early development and allometric growth in hatchery-reared Eurasian perch, Perca fluviatilis L. Aquac Res 50. 10.1111/are.14208

Langfelder, P., Horvath, S., 2008. WGCNA: An R package for weighted correlation network analysis. BMC Bioinformatics 9. 10.1186/1471-2105-9-559

Leesch, F., Lorenzo-Orts, L., Pribitzer, C., Grishkovskaya, I., Roehsner, J., Chugunova, A., Matzinger, M., Roitinger, E., Belačić, K., Kandolf, S., Lin, T.Y., Mechtler, K., Meinhart, A., Haselbach, D., Pauli, A., 2023. A molecular network of conserved factors keeps ribosomes dormant in the egg. Nature 613. 10.1038/s41586-022-05623-y

Love, M.I., Huber, W., Anders, S., 2014. Moderated estimation of fold change and dispersion for RNA-seq data with DESeq2. Genome Biol 15. 10.1186/s13059-014-0550-8

Mathavan, S., Lee, S.G.P., Mak, A., Miller, L.D., Murthy, K.R.K., Govindarajan, K.R., Tong, Y., Wu, Y.L., Lam, S.H., Yang, H., Ruan, Y., Korzh, V., Gong, Z., Liu, E.T., Lufkin, T., 2005. Transcriptome analysis of zebrafish embryogenesis using microarrays. PLoS Genet 1. 10.1371/journal.pgen.0010029

Mazurais, D., Darias, M., Zambonino-Infante, J.L., Cahu, C.L., 2011. Transcriptomics for understanding marine fish larval development. Can J Zool. 10.1139/z11-036

McMenamin, S.K., Parichy, D.M., 2013. Chapter Five – Metamorphosis in Teleosts, in: Current Topics in Developmental Biology.

Migaud, H., Bell, G., Cabrita, E., Mcandrew, B., Davie, A., Bobe, J., Herráez, M.P., Carrillo, M., 2013. Gamete quality and broodstock management in temperate fish. Rev Aquac. 10.1111/raq.12025

Nelson, J.C., Granato, M., 2022. Zebrafish behavior as a gateway to nervous system assembly and plasticity. Development (Cambridge) 149. 10.1242/dev.177998

Nynca, J., Ciereszko, A., 2009. Measurement of concentration and viability of brook trout (Salvelinus fontinalis)spermatozoa using computer-aided fluorescent microscopy. Aquaculture 292. 10.1016/j.aquaculture.2009.04.020

Osse, J.W.M., Van Den Boogaart, J.G.M., Van Snik, G.M.J., Van Der Sluys, L., 1997. Priorities during early growth of fish larvae, in: Aquaculture. 10.1016/S0044-8486(97)00126-9

Palińska-Żarska, K., Król, J., Woźny, M., Kamaszewski, M., Szudrowicz, H., Wiechetek, W., Brzuzan, P., Fopp-Bayat, D., Żarski, D., 2021. Domestication affected stress and immune response markers in Perca fluviatilis in the early larval stage. Fish Shellfish Immunol 114. 10.1016/j.fsi.2021.04.028

Palińska-Żarska, K., Woźny, M., Kamaszewski, M., Szudrowicz, H., Brzuzan, P., Żarski, D., 2020. Domestication process modifies digestion ability in larvae of Eurasian perch (Perca fluviatilis), a freshwater Teleostei. Sci Rep 10. 10.1038/s41598-020-59145-6

Pertea, M., Pertea, G.M., Antonescu, C.M., Chang, T.C., Mendell, J.T., Salzberg, S.L., 2015. StringTie enables improved reconstruction of a transcriptome from RNA-seq reads. Nat Biotechnol 33. 10.1038/nbt.3122

Policar, T., Schaefer, F.J., Panana, E., Meyer, S., Teerlinck, S., Toner, D., Żarski, D., 2019. Recent progress in European percid fish culture production technology—tackling bottlenecks. Aquaculture International. 10.1007/s10499-019-00433-y

Qi, S.T., Ma, J.Y., Wang, Z.B., Guo, L., Hou, Y., Sun, Q.Y., 2016. N6 -methyladenosine sequencing highlights the involvement of mRNA methylation in oocyte meiotic maturation and embryo development by regulating translation in xenopus laevis. Journal of Biological Chemistry 291. 10.1074/jbc.M116.748889

Romano, A., Barca, A., Storelli, C., Verri, T., 2014. Teleost fish models in membrane transport research: The PEPT1(SLC15A1) H+-oligopeptide transporter as a case study. Journal of Physiology 592. 10.1113/jphysiol.2013.259622

Sarosiek, B., Dryl, K., Krejszeff, S., Zarski, D., 2016. Characterization of pikeperch (Sander lucioperca) milt collected with a syringe and a catheter. Aquaculture 450. 10.1016/j.aquaculture.2015.06.040

Sarropoulou, E., Tsalafouta, A., Sundaram, A.Y.M., Gilfillan, G.D., Kotoulas, G., Papandroulakis, N., Pavlidis, M., 2016. Transcriptomic changes in relation to early-life events in the gilthead sea bream (Sparus aurata). BMC Genomics 17. 10.1186/s12864-016-2874-0

Sayers, E.W., Bolton, E.E., Brister, J.R., Canese, K., Chan, J., Comeau, D.C., Connor, R., Funk, K., Kelly, C., Kim, S., Madej, T., Marchler-Bauer, A., Lanczycki, C., Lathrop, S., Lu, Z., Thibaud-Nissen, F., Murphy, T., Phan, L., Skripchenko, Y., Tse, T., Wang, J., Williams, R., Trawick, B.W., Pruitt, K.D., Sherry, S.T., 2022. Database resources of the national center for biotechnology information. Nucleic Acids Res 50. 10.1093/nar/gkab1112

Schmittgen, T.D., Livak, K.J., 2001. Analysis of relative gene expression data using real-time quantitative PCR and the 2(-Delta Delta C(T)) Method. Methods 25.

Shen, Z.G., Yao, H., Guo, L., Li, X.X., Wang, H.P., 2017. Ribosome RNA Profiling to Quantify Ovarian Development and Identify Sex in Fish. Sci Rep 7. 10.1038/s41598-017-04327-y

Simon Andrews, 2020. Babraham Bioinformatics - FastQC A Quality Control tool for High Throughput Sequence Data. Soil.

Subasinghe, R., Soto, D., Jia, J., 2009. Global aquaculture and its role in sustainable development. Rev Aquac 1. 10.1111/j.1753-5131.2008.01002.x

Sumana, S.L., Chen, H., Shui, Y., Zhang, C., Yu, F., Zhu, J., Su, S., 2023. Effect of Dietary Selenium on the Growth and Immune Systems of Fish. Animals. 10.3390/ani13182978

Team, R.C., 2021. R: A Language and Environment for Statistical Computing. R Foundation for Statistical Computing.

Toomey, L., Dellicour, S., Kapusta, A., Żarski, D., Buhrke, F., Milla, S., Fontaine, P., Lecocq, T., 2021. Split it up and see: using proxies to highlight divergent inter-populational performances in aquaculture standardised conditions. BMC Ecol Evol 21. 10.1186/s12862-021-01937-z

Untergasser, A., Nijveen, H., Rao, X., Bisseling, T., Geurts, R., Leunissen, J.A.M., 2007. Primer3Plus, an enhanced web interface to Primer3. [Nucleic Acids Res. 2007] - PubMed result. Nucleic Acids Res 35.

Urho, L., 2002. Characters of larvae - What are they? Folia Zool Brno.

Vacca, F., Barca, A., Gomes, A.S., Mazzei, A., Piccinni, B., Cinquetti, R., Del Vecchio, G., Romano, A., Rønnestad, I., Bossi, E., Verri, T., 2019. The peptide transporter 1a of the zebrafish Danio rerio, an emerging model in nutrigenomics and nutrition research: Molecular characterization, functional properties, and expression analysis. Genes Nutr 14. 10.1186/s12263-019-0657-3

Valente, L.M.P., Moutou, K.A., Conceição, L.E.C., Engrola, S., Fernandes, J.M.O., Johnston, I.A., 2013. What determines growth potential and juvenile quality of farmed fish species? Rev Aquac 5. 10.1111/raq.12020

van der Aa, L.M., Levraud, J.P., Yahmi, M., Lauret, E., Briolat, V., Herbomel, P., Benmansour, A., Boudinot, P., 2009. A large new subset of TRIM genes highly diversified by duplication and positive selection in teleost fish. BMC Biol 7. 10.1186/1741-7007-7-7

Wei, T., Simko, V., Levy, M., Xie, Y., Jin, Y.J., Zemla, J., 2017. Visualization of a Correlation Matrix. R package “corrplot”. Statistician 56.

Yúfera, M., Darias, M.J., 2007. The onset of exogenous feeding in marine fish larvae. Aquaculture. 10.1016/j.aquaculture.2007.04.050

Żarski, D., Ben Ammar, I., Bernáth, G., Baekelandt, S., Bokor, Z., Palińska-Żarska, K., Fontaine, P., Horváth, Á., Kestemont, P., Mandiki, S.N.M., 2020a. Repeated hormonal induction of spermiation affects the stress but not the immune response in pikeperch (Sander lucioperca). Fish Shellfish Immunol 101. 10.1016/j.fsi.2020.03.057

Zarski, D., Bokor, Z., Kotrik, L., Urbanyi, B., Horváth, A., Targońska, K., Krejszeff, S., Palińska, K., Kucharczyk, D., 2011. A new classification of a preovulatory oocyte maturation stage suitable for the synchronization of ovulation in controlled reproduction of Eurasian perch, Perca fluviatilis L. Reprod Biol 11. 10.1016/S1642-431X(12)60066-7

Żarski, D., Horváth, Á., Bernáth, G., Krejszeff, S., Radóczi, J., Palińska-Żarska, K., Bokor, Z., Kupren, K., Urbányi, B., 2017a. Harvest, Transport of Spawners, Prophylaxis. 10.1007/978-3-319-49376-3_2

Żarski, D., Horváth, Á., Bernáth, G., Krejszeff, S., Radóczi, J., Palińska-Żarska, K., Bokor, Z., Kupren, K., Urbányi, B., 2017b. Collection of Gametes. 10.1007/978-3-319-49376-3_6

Żarski, D., Horváth, Á., Bernáth, G., Krejszeff, S., Radóczi, J., Palińska-Żarska, K., Bokor, Z., Kupren, K., Urbányi, B., 2017c. In Vitro Fertilization. 10.1007/978-3-319-49376-3_9

Żarski, D., Horváth, Á., Bernáth, G., Krejszeff, S., Radóczi, J., Palińska-Żarska, K., Bokor, Z., Kupren, K., Urbányi, B., 2017d. Incubation and Hatching. 10.1007/978-3-319-49376-3_10

Żarski, D., Le Cam, A., Frohlich, T., Kösters, M., Klopp, C., Nynca, J., Ciesielski, S., Sarosiek, B., Dryl, K., Montfort, J., Król, J., Fontaine, P., Ciereszko, A., Bobe, J., 2021. Neurodevelopment vs. the immune system: Complementary contributions of maternally-inherited gene transcripts and proteins to successful embryonic development in fish. Genomics 113. 10.1016/j.ygeno.2021.09.003

Żarski, D., Le Cam, A., Nynca, J., Klopp, C., Ciesielski, S., Sarosiek, B., Montfort, J., Król, J., Fontaine, P., Ciereszko, A., Bobe, J., 2020b. Domestication modulates the expression of genes involved in neurogenesis in high-quality eggs of Sander lucioperca. Mol Reprod Dev 87. 10.1002/mrd.23414

Żarski, D., Nguyen, T., Le Cam, A., Montfort, J., Dutto, G., Vidal, M.O., Fauvel, C., Bobe, J., 2017e. Transcriptomic Profiling of Egg Quality in Sea Bass (Dicentrarchus labrax) Sheds Light on Genes Involved in Ubiquitination and Translation. Marine Biotechnology 19. 10.1007/s10126-017-9732-1

Zhao, S., Fernald, R.D., 2005. Comprehensive algorithm for quantitative real-time polymerase chain reaction. Journal of Computational Biology 12. 10.1089/cmb.2005.12.1047

