## Supplementary file 1 for "Does transcriptome of freshly hatched fish larvae describe past or predict future developmental trajectory?"

**Supplementary file S1 :** Broodstock characteristics

| FAMILIES | BROODSTOCK | ORIGIN | WEIGHT (g) | TOTAL LENGTH (cm) | FORK LENGTH (cm) | AGE |
|---|---|---|---|---|---|---|
| **F1** | FEMALE 1 | Żurawia | 343 | 28.2 | 24.8 | 6 |
| | MALE 1 | Żurawia(Rytwiany Pond System) | 203 | 25.7 | 21.3 | 4 |
| **F2** | FEMALE 2 | Szymon lake | 571 | 32 | 27.5 | 7 |
| | MALE 2 | Żurawia(Rytwiany Pond System ) | 359 | 29.5 | 26 | 4 |
| **F3** | FEMALE 3 | Żurawia | 236 | 24.6 | 21.8 | 5 |
| | MALE 3 | Żurawia(Rytwiany Pond System) | 151 | 23.2 | 20 | 4 |
| **F4** | FEMALE 4 | Żurawia | 299 | 26.8 | 23.8 | 5 |
| | MALE 4 | Żurawia(Rytwiany Pond System) | 242 | 25.9 | 22.8 | 5 |
| **F5** | FEMALE 5 | Żurawia | 302 | 25.8 | 24 | 6 |
| | MALE 5 | Żurawia(Rytwiany Pond System) | 152 | 23.5 | 20.5 | 4+ |
| **F6** | FEMALE 6 | Żurawia | 297 | 25 | 24 | 6 |
| | MALE 6 | Żurawia(Rytwiany Pond System) | 218 | 25.9 | 23 | 5+ |
| **F7** | FEMALE 7 | Umląg lake | 216 | 23.5 | 21.5 | 5 |
| | MALE 7 | Żurawia(Rytwiany Pond System) | 154 | 23.5 | 20.3 | ? |
| **F8** | FEMALE 8 | Żurawia | 285 | 26.5 | 23 | 5 |
| | MALE 8 | Żurawia(Rytwiany Pond System) | 196 | 24.6 | 21.5 | 5+ |
| **F9** | FEMALE 9 | Szymon lake | 608 | 35.5 | 32 | 7 |
| | MALE 9 | Żurawia(Rytwiany Pond System) | 196 | 25.5 | 22.5 | ? |
| **F10** | FEMALE 10 | Szymon lake | 763 | 35 | 31.4 | 8 |
| | MALE 10 | Żurawia(Rytwiany Pond System) | 160 | 24.1 | 20.8 | ? |
| **F11** | FEMALE 11 | Umląg lake | 336 | 28 | 24.5 | 6 |
| | MALE 11 | Żurawia(Rytwiany Pond System) | 137 | 22 | 19.2 | ? |
| **F12** | FEMALE 12 | Umląg lake | 810 | 36.5 | 33 | 7 |
| | MALE 12 | Żurawia(Rytwiany Pond System) | 105 | 20.7 | 17.5 | ? |
| **F13** | FEMALE 13 | Szymon lake | 611 | 35.3 | 30.8 | 7 |
| | MALE 13 | Żurawia(Rytwiany Pond System) | 142 | 22.5 | 19.8 | ? |
| **F14** | FEMALE 14 | Szymon lake | 725 | 37.2 | 32 | 7 |
| | MALE 14 | Iława | 246 | 26.1 | 23 | 4 |
| **F15** | FEMALE 15 | Szymon lake | 720 | 37.3 | 32 | 5 |
| | MALE 15 | Iława | 223 | 25.8 | 22.2 | 4 |
| **F16** | FEMALE 16 | Szymon lake | 695 | 34.3 | 30.5 | 6 |
| | MALE 16 | Żurawia(Rytwiany Pond System) | 133 | 22.2 | 18.5 | 3+ |
