## Supplementary file 2 for "Does transcriptome of freshly hatched fish larvae describe past or predict future developmental trajectory?"

**Supplementary file S2**: Sperm evaluation with CASA before and after cryopreservation. ALH: Amplitude of Lateral Head Displacement; LIN: Linearity; VAP: Average Path Velocity; VCL: Curvilinear Velocity; VSl: Straight Velocity; MOT: percentage of sperm motility

| Males | ALH (μm) | LIN (%) | VAP (μm $s^{-1}$) | VCL (μm $s^{-1}$) | VSL (μm $s^{-1}$) | MOT(%) |
|---|---|---|---|---|---|---|
| 1 | 7.556148 | 72.14422 | 184.0915 | 205.7676 | 163.0863 | 75.3 |
| 2 | 7.933224 | 81.7436 | 227.7508 | 243.9181 | 201.3321 | 87.6 |
| 3 | 8.887574 | 81.3074 | 228.7303 | 245.4705 | 202.2623 | 80.4 |
| 4 | 10.35598 | 75.90818 | 214.9773 | 236.3426 | 182.0324 | 90 |
| 5 | 7.100678 | 74.39593 | 175.9103 | 198.9887 | 153.0129 | 82.7 |
| 6 | 6.693855 | 79.79078 | 183.4773 | 200.7846 | 164.2914 | 84.2 |
| 7 | 7.257437 | 80.20573 | 197.5499 | 211.8014 | 173.8595 | 80.2 |
| 8 | 8.525166 | 74.15175 | 191.4987 | 214.5812 | 162.5481 | 90.5 |
| 9 | 7.811111 | 79.93286 | 199.2677 | 216.7182 | 176.257 | 86.9 |
| 10 | 7.653057 | 75.85913 | 183.2043 | 205.2517 | 157.7023 | 89.3 |
| 11 | 6.946595 | 78.89905 | 190.2624 | 208.2974 | 167.1579 | 86.1 |
| 12 | 6.497107 | 82.37009 | 194.6677 | 209.386 | 176.4434 | 80.9 |
| 13 | 8.169308 | 77.52857 | 194.3661 | 214.7507 | 168.3114 | 89.7 |
| 14 | 7.514402 | 77.69066 | 189.9809 | 210.0948 | 166.0552 | 90.3 |
| 15 | 7.065888 | 77.29461 | 182.6261 | 203.5702 | 158.9418 | 87.9 |
| 16 | 8.346438 | 78.78947 | 200.7829 | 219.6534 | 174.7468 | 85.7 |
| 1 cryo | 5.183588 | 67.98289 | 117.5497 | 145.9357 | 104.5426 | 61.3 |
| 2 cryo | 4.540228 | 70.2931 | 120.6054 | 147.0256 | 112.78 | 50.8 |
| 3 cryo | 5.096318 | 68.08394 | 120.8708 | 151.0514 | 110.6403 | 58.6 |
| 4 cryo | 4.124404 | 64.94624 | 95.60346 | 126.8639 | 86.79023 | 63.2 |
| 5 cryo | 5.029983 | 65.09456 | 118.4535 | 147.959 | 104.9009 | 59.8 |
| 6 cryo | 4.504754 | 63.51034 | 103.8411 | 134.6898 | 94.96765 | 55 |
| 7 cryo | 4.495647 | 61.96903 | 100.6436 | 128.4414 | 92.18875 | 52.1 |
| 8 cryo | 4.389957 | 64.37805 | 104.8808 | 134.6037 | 96.41796 | 56.3 |
| 9 cryo | 4.449642 | 62.10599 | 98.48766 | 130.6613 | 89.44179 | 59.6 |
| 10 cryo | 4.273771 | 72.05066 | 119.7826 | 145.1478 | 111.0787 | 59.8 |
| 11 cryo | 4.425791 | 59.93989 | 91.90695 | 127.2347 | 83.11011 | 59.6 |
| 12 cryo | 4.201456 | 73.01287 | 123.2177 | 146.9202 | 114.8439 | 57.4 |
| 13 cryo | 4.267513 | 63.86021 | 101.0722 | 132.3957 | 92.65259 | 54.2 |
| 14 cryo | 4.409236 | 65.01599 | 108.3475 | 132.5867 | 98.41082 | 75 |
| 15 cryo | 4.547335 | 69.00627 | 113.5295 | 137.3622 | 102.7892 | 74.4 |
| 16 cryo | 4.926946 | 75.25546 | 144.1598 | 166.9635 | 130.6906 | 73.2 |
