## Supplementary file 3 for "Does transcriptome of freshly hatched fish larvae describe past or predict future developmental trajectory?"

**Supplementary file S3:** Genes with high correlations (both positive and negative) to specific Key aquaculture traits. A. Genes related to cannibalism. B. Genes highly correlated to mortality C. Genes most correlated to SBIE. D. Genes highly correlated to SGR for length E. Genes highly correlated to SGR for weight. F. Genes highly correlated to weight of larvae at the end of the experiment G. Genes highly correlated to Fulton's condition factor. GS: Gene Significance

## A.

| GENE_ID | TRANSCRIPT_ACCESSION_NUMBER | HUMAN ORTHOLOGS | GS | *p-value* |
|---|---|---|---|---|
| *selenoo2* | XM_039791163.1 | *SELENOO* | 0.823647 | 8.80E-05 |
| *pex2* | XM_039790640.1 | *PEX2* | 0.759129 | 0.000649 |
| *LOC120559301* | XM_039800928.1 | *TMEM198* | 0.748497 | 0.000851 |
| *LOC120558849* | XM_039800144.1 | *NA* | 0.744127 | 0.000948 |
| *LOC120560490* | XM_039803062.1 | *ABCB9* | 0.722085 | 0.001585 |
| *LOC120543986* | XM_039777447.1 | *AR* | 0.711043 | 0.002015 |
| *nudt12* | XM_039803805.1 | *NUDT12* | -0.84789 | 3.35E-05 |
| *LOC120573883* | XM_039823776.1 | *NA* | -0.83261 | 6.27E-05 |
| *LOC120547285* | XM_039782855.1 | *MTUS1* | -0.78394 | 0.000326 |
| *LOC120556999* | XM_039796989.1 | *TASOR* | -0.78027 | 0.000363 |
| *mmp25b* | XM_039788073.1 | *MMP17* | -0.77709 | 0.000397 |
| *c8h15orf40* | XM_039807462.1 | *C15orf40* | -0.76682 | 0.000529 |
| *pds5b* | XM_039776878.1 | *PDS5B* | -0.76502 | 0.000555 |
| *rhbdl1* | XM_039787716.1 | *RHBDL1* | -0.76073 | 0.000622 |
| *sh2d3cb* | XM_039802731.1 | *SH2D3C* | -0.75116 | 0.000796 |
| *cmah* | XM_039791006.1 | *CMAHP* | -0.74459 | 0.000937 |
| *LOC120549684* | XM_039786766.1 | *NA* | -0.73278 | 0.001243 |
| *LOC120568916* | XM_039816668.1 | *MARCHF2* | -0.72786 | 0.001391 |
| *LOC120571615* | XR_005641248.1 | *NA* | -0.72719 | 0.001413 |
| *lactbl1b* | XM_039798927.1 | *LACTBL1* | -0.72094 | 0.001626 |
| *nid2a* | XM_039793428.1 | *NID2* | -0.7084 | 0.002131 |
| *rxfp3.2b* | XM_039810374.1 | *RXFP3* | -0.70289 | 0.002389 |

## B.

| GENE_ID | TRANSCRIPT_ACCESSION_NUMBER | HUMAN ORTHOLOGS | GS | *p-value* |
|---|---|---|---|---|
| *mkxa* | XM_039789540.1 | *MKX* | 0.773396 | 0.000441 |
| *LOC120557731* | XM_039798307.1 | *TCP11L1* | 0.771872 | 0.00046 |
| *LOC120575215* | XM_039825900.1 | *SOX8* | 0.757658 | 0.000674 |
| *LOC120554598* | XM_039793599.1 | *IL1RAPL1* | 0.754489 | 0.000732 |
| *rtn4rl2a* | XM_039816981.1 | *RTN4RL2* | 0.753633 | 0.000748 |
| *LOC120556829* | XM_039796566.1 | *CPNE1* | 0.739755 | 0.001054 |
| *klf5a* | XM_039782934.1 | *KLF5* | 0.739543 | 0.001059 |
| *fynb* | XM_039782603.1 | *FYN* | 0.738856 | 0.001077 |
| *LOC120571709* | XM_039820779.1 | *CSPG5* | 0.725929 | 0.001454 |
| *si:ch1073-145m9.1* | XM_039821276.1 | *NA* | 0.725098 | 0.001481 |

| | | | | |
|---|---|---|---|---|
| *LOC120574026* | XM_039824018.1 | *PDZK1* | 0.723493 | 0.001536 |
| *LOC120557984* | XM_039798751.1 | *BHLHE40* | 0.719774 | 0.001668 |
| *morn4* | XM_039784652.1 | *MORN4* | 0.719557 | 0.001676 |
| *rcbtb1* | XM_039793148.1 | *RCBTB1* | 0.718855 | 0.001702 |
| *pdcd4a* | XM_039797508.1 | *PDCD4* | 0.718174 | 0.001728 |
| *lnx2a* | XM_039790511.1 | *LNX2* | 0.713051 | 0.00193 |
| *bmpr1ba* | XM_039804081.1 | *BMPR1B* | 0.711197 | 0.002008 |
| *LOC120551619* | XM_039789114.1 | *KCNH7* | 0.710976 | 0.002018 |
| *si:ch211-215i13.3* | XM_039819213.1 | *BAALC* | 0.706731 | 0.002206 |
| *rapgef2* | XM_039813634.1 | *RAPGEF2* | 0.705271 | 0.002275 |
| *zgc:122979* | XM_039778904.1 | *DNAJB5* | 0.704972 | 0.002289 |
| *LOC120570925* | XM_039819623.1 | *HLF* | 0.703696 | 0.00235 |
| *fam43b* | XM_039799236.1 | *FAM43B* | 0.70188 | 0.00244 |
| *LOC120575713* | XM_039826547.1 | *MLXIPL* | 0.700779 | 0.002495 |
| *LOC120544023* | XM_039777522.1 | *CRCP* | -0.80575 | 0.000165 |
| *cnpy2* | XM_039798701.1 | *CNPY2* | -0.77553 | 0.000415 |
| *si:ch211-217a12.1* | XM_039790223.1 | *GPT2* | -0.76661 | 0.000532 |
| *adck2* | XM_039808041.1 | *ADCK2* | -0.75944 | 0.000644 |
| *LOC120567770* | XM_039814785.1 | *GTF3A* | -0.7538 | 0.000745 |
| *polr1f* | XM_039821120.1 | *POLR1F* | -0.7476 | 0.00087 |
| *cxcr3.2* | XM_039805568.1 | *CXCR2* | -0.74109 | 0.001021 |
| *LOC120563727* | XM_039808101.1 | *ERGIC2* | -0.73826 | 0.001092 |
| *enoph1* | XM_039780775.1 | *ENOPH1* | -0.73681 | 0.00113 |
| *gmpr2* | XM_039821768.1 | *GMPR2* | -0.72868 | 0.001366 |
| *ddx52* | XM_039778580.1 | *DDX52* | -0.72852 | 0.001371 |
| *LOC120555473* | XM_039794172.1 | *PLEKHF2* | -0.72695 | 0.001421 |
| *LOC120558123* | XM_039799029.1 | *DTNBP1* | -0.72347 | 0.001537 |
| *dnajc30b* | XM_039786891.1 | *DNAJC30* | -0.7209 | 0.001627 |
| *uts2d* | XM_039812033.1 | *XOXO* | -0.72076 | 0.001632 |
| *LOC120546050* | XM_039780806.1 | *STOML2* | -0.71954 | 0.001677 |
| *si:ch211-214j24.10* | XM_039792521.1 | *NA* | -0.71752 | 0.001753 |
| *pus3* | XM_039788994.1 | *PUS3* | -0.71697 | 0.001773 |
| *tomm20a* | XM_039784734.1 | *TOMM20* | -0.71547 | 0.001832 |
| *mpp6b* | XM_039820817.1 | *PALS2* | -0.71288 | 0.001937 |
| *LOC120545887* | XM_039780516.1 | *PBDC1* | -0.71155 | 0.001993 |
| *golt1bb* | XM_039807427.1 | *GOLT1B* | -0.71108 | 0.002013 |
| *lactb* | XM_039808237.1 | *LACTB* | -0.71033 | 0.002045 |
| *LOC120561432* | XM_039804553.1 | *TTF1* | -0.70951 | 0.002081 |
| *cacng5b* | XM_039823832.1 | *CACNG5* | -0.70509 | 0.002283 |
| *LOC120573711* | XM_039823626.1 | *CAAP1* | -0.70418 | 0.002327 |

## C.

| GENE_ID | TRANSCRIPT_ACCESSION_NUMBER | HUMAN ORTHOLOGS | GS | *p-value* |
|---|---|---|---|---|

| | | | | |
|---|---|---|---|---|
| *si:dkey-117m1.4* | XM_039803324.1 | *NA* | 0.791894 | 0.000256 |
| *arsg* | XM_039788329.1 | *ARSG* | 0.769645 | 0.000489 |
| *kdrl* | XM_039813298.1 | *FLT1* | 0.738975 | 0.001074 |
| *LOC120561638* | XM_039804800.1 | *NA* | 0.737386 | 0.001115 |
| *malsu1* | XM_039807250.1 | *MALSU1* | 0.725822 | 0.001457 |
| *npdc1b* | XM_039802865.1 | *NPDC1* | 0.70592 | 0.002244 |
| *barx2* | XM_039821365.1 | *BARX2* | 0.701486 | 0.002459 |
| *LOC120575397* | XM_039826188.1 | *PYCARD* | -0.77819 | 0.000385 |
| *aggf1* | XM_039797527.1 | *AGGF1* | -0.76316 | 0.000583 |
| *LOC120567195* | XM_039814111.1 | *PCDHGC5* | -0.75051 | 0.000809 |
| *cubn* | XM_039789731.1 | *CUBN* | -0.73351 | 0.001222 |
| *lrrk1* | XM_039794699.1 | *LRRK1* | -0.73125 | 0.001287 |
| *tbc1d30* | XM_039792075.1 | *TBC1D30* | -0.72395 | 0.00152 |
| *st7l* | XM_039798844.1 | *ST7* | -0.71834 | 0.001721 |
| *tmem216* | XM_039813676.1 | *TMEM216* | -0.71455 | 0.001869 |
| *LOC120556874* | XM_039796687.1 | *NA* | -0.70092 | 0.002488 |

**D.**

| **GENE_ID** | **TRANSCRIPT_ACCESSION_NUMBER** | **HUMAN ORTHOLOGS** | **GS** | ***p-value*** |
|---|---|---|---|---|
| *atp8a2* | XM_039789439.1 | *ATP8A2* | 0.873817 | 9.72E-06 |
| *gpr179* | XM_039803243.1 | *GPR158* | 0.764197 | 0.000567 |
| *LOC120571047* | XM_039819747.1 | *ZNF184* | 0.758972 | 0.000651 |
| *polr2b* | XM_039813488.1 | *POLR2B* | 0.753756 | 0.000745 |
| *dscama* | XM_039777637.1 | *DSCAM* | 0.753162 | 0.000757 |
| *tekt1* | XM_039820506.1 | *TEKT1* | 0.747324 | 0.000876 |
| *adamtsl7* | XM_039801823.1 | *THSD4* | 0.742693 | 0.000982 |
| *gpr176* | XM_039782698.1 | *GPR176* | 0.730583 | 0.001307 |
| *chrnb1* | XM_039823570.1 | *CHRNB1* | 0.721703 | 0.001598 |
| *plxnc1* | XM_039792449.1 | *PLXNC1* | 0.70922 | 0.002094 |
| *LOC120550477* | XM_039787166.1 | *A2ML1* | 0.707241 | 0.002183 |
| *slc15a1a* | XM_039817133.1 | *SLC15A1* | -0.8001 | 0.000198 |
| *cln3* | XM_039825647.1 | *CLN3* | -0.79474 | 0.000235 |
| *LOC120547353* | XR_005637056.1 | *NA* | -0.79011 | 0.000271 |
| *faxca* | XM_039782711.1 | *FAXC* | -0.76895 | 0.000499 |
| *pnp5a* | XM_039823439.1 | *PNP* | -0.76177 | 0.000605 |
| *cuedc2* | XM_039783959.1 | *CUEDC2* | -0.74137 | 0.001014 |
| *LOC120549428* | XM_039786345.1 | *THBS1* | -0.7346 | 0.001191 |
| *LOC120567821* | XM_039814879.1 | *MYOC* | -0.73376 | 0.001215 |
| *loxl3b* | XM_039785199.1 | *LOXL3* | -0.72527 | 0.001476 |
| *LOC120559460* | XM_039801160.1 | *NKX2-3* | -0.72465 | 0.001496 |
| *ccbe1* | XM_039780041.1 | *CCBE1* | -0.72083 | 0.001629 |
| *LOC120565036* | XM_039810342.1 | *ALPI* | -0.71272 | 0.001944 |
| *fam221a* | XM_039819369.1 | *FAM221A* | -0.71005 | 0.002058 |
| *elna* | XM_039824656.1 | *NA* | -0.70996 | 0.002062 |
| *LOC120554227* | XM_039792984.1 | *ZNF84* | -0.70834 | 0.002133 |
| *ythdc1* | XM_039779087.1 | *YTHDC1* | -0.70611 | 0.002235 |

| | | | | |
|---|---|---|---|---|
| *LOC120559933* | XR_005639388.1 | *XOXO* | -0.70561 | 0.002259 |
| *LOC120570652* | XM_039819131.1 | *PER2* | -0.70494 | 0.00229 |
| *si:ch211-176g6.2* | XM_039789392.1 | *RADIL* | -0.70441 | 0.002316 |
| *LOC120549424* | XM_039786332.1 | *CHAC1* | -0.70326 | 0.002371 |
| *csnk2a2b* | XM_039809336.1 | *CSNK2A2* | -0.70322 | 0.002373 |
| *gareml* | XM_039782726.1 | *GAREM2* | -0.70121 | 0.002473 |

**E.**

| **GENE_ID** | **TRANSCRIPT_ACCESSION_NUMBER** | **HUMAN ORTHOLOGS** | **GS** | ***p-value*** |
|---|---|---|---|---|
| *prdm1b* | XM_039821118.1 | *PRDM1* | 0.83547 | 5.60E-05 |
| *agpat4* | XM_039783933.1 | *AGPAT4* | 0.780512 | 0.00036 |
| *apoba* | XM_039785565.1 | *APOB* | 0.778804 | 0.000378 |
| *tgfbr3* | XM_039810773.1 | *TGFBR3* | 0.766218 | 0.000537 |
| *LOC120546953* | XM_039782234.1 | *EVA1C* | 0.756307 | 0.000698 |
| *ildr2* | XM_039818909.1 | *ILDR2* | 0.750336 | 0.000813 |
| *dtx4a* | XM_039823160.1 | *DTX4* | 0.739289 | 0.001066 |
| *c20h14orf180* | XM_039785648.1 | *CNST* | 0.731303 | 0.001286 |
| *zgc:171971* | XM_039788201.1 | *POLR3D* | 0.730263 | 0.001317 |
| *LOC120568699* | XM_039816370.1 | *SLC25A24* | 0.729841 | 0.00133 |
| *neurl1aa* | XM_039776668.1 | *NEURL1* | 0.722336 | 0.001576 |
| *arrb1* | XM_039777112.1 | *ARRB1* | 0.717657 | 0.001747 |
| *LOC120564302* | XM_039809154.1 | *H1-4* | 0.716955 | 0.001774 |
| *LOC120545456* | XM_039779784.1 | *CSPG4* | 0.713425 | 0.001915 |
| *LOC120559901* | XM_039801983.1 | *TRIM7* | 0.712669 | 0.001946 |
| *slc4a5b* | XM_039779149.1 | *SLC4A5* | 0.711218 | 0.002007 |
| *LOC120555597* | XM_039794414.1 | *B3GALT2* | 0.706184 | 0.002232 |
| *LOC120564168* | XM_039808937.1 | *ARL2BP* | 0.704097 | 0.002331 |
| *tfap2a* | XM_039790007.1 | *TFAP2A* | 0.703349 | 0.002367 |
| *txnl4a* | XM_039821238.1 | *TXNL4A* | -0.84056 | 4.56E-05 |
| *muc13b* | XM_039789342.1 | *MUC13B* | -0.80523 | 0.000167 |
| *mid1ip1l* | XM_039813944.1 | *MID1IP1* | -0.80393 | 0.000175 |
| *c21h10orf88* | XM_039789081.1 | *PAAT* | -0.75839 | 0.000661 |
| *si:ch211-59o9.10* | XM_039807797.1 | *RNF38* | -0.75607 | 0.000703 |
| *nkapd1* | XM_039782424.1 | *NKAPD1* | -0.74988 | 0.000822 |
| *LOC120547184* | XM_039782660.1 | *KCNK2* | -0.74846 | 0.000852 |
| *zanl* | XM_039823009.1 | *ZAN* | -0.74358 | 0.000961 |
| *si:dkey-57a22.11* | XM_039818970.1 | *SGO1* | -0.74333 | 0.000967 |
| *LOC120557765* | XM_039798369.1 | *NAT8* | -0.73551 | 0.001166 |
| *LOC120544654* | XM_039778556.1 | *RRM1* | -0.73228 | 0.001257 |
| *fabp2* | XM_039806486.1 | *FABP2* | -0.72505 | 0.001483 |
| *chtf8* | XM_039808917.1 | *CHTF8* | -0.72309 | 0.00155 |
| *LOC120556667* | XM_039796312.1 | *CHIA* | -0.72078 | 0.001631 |
| *LOC120544521* | XM_039778329.1 | *CIPC* | -0.71893 | 0.001699 |
| *cdc14aa* | XM_039816763.1 | *CDC14A* | -0.71262 | 0.001948 |

| | | | | |
|---|---|---|---|---|
| *LOC120562327* | XM_039806016.1 | *ZNF154* | -0.70764 | 0.002165 |
| *cryz* | XM_039815914.1 | *CRYZ* | -0.70485 | 0.002295 |
| *mrps6* | XM_039784102.1 | *MRPS6* | -0.70438 | 0.002317 |
| *tango2* | XM_039804096.1 | *TANGO2* | -0.70236 | 0.002415 |
| *cald1b* | XM_039808207.1 | *CALD1* | -0.70128 | 0.00247 |

**F.**

| **GENE_ID** | **TRANSCRIPT_ACCESSION_NUMBER** | **HUMAN ORTHOLOGS** | **GS** | ***p-value*** |
|---|---|---|---|---|
| *LOC120552004* | XM_039789614.1 | *AOC1* | 0.811459 | 0.000136 |
| *gpr176* | XM_039782698.1 | *GPR176* | 0.778565 | 0.000381 |
| *pou2f3* | XM_039826492.1 | *POU2F3* | 0.768447 | 0.000506 |
| *hck* | XM_039800624.1 | *HCK* | 0.764008 | 0.00057 |
| *LOC120555597* | XM_039794414.1 | *B3GALT2* | 0.75433 | 0.000735 |
| *LOC120550477* | XM_039787166.1 | *A2ML1* | 0.747012 | 0.000883 |
| *LOC120553155* | XM_039791254.1 | *PAH* | 0.743519 | 0.000962 |
| *ddc* | XM_039806765.1 | *DDC* | 0.737704 | 0.001107 |
| *ccser1* | XM_039802935.1 | *CCSER1* | 0.72681 | 0.001425 |
| *b3gat2* | XM_039784296.1 | *B3GAT2* | 0.71187 | 0.00198 |
| *LOC120564637* | XM_039809793.1 | *KBTBD13* | 0.70476 | 0.002299 |
| *fam167ab* | XM_039781499.1 | *FAM167A* | 0.704741 | 0.0023 |
| *rhot1b* | XM_039805836.1 | *RHOT1* | 0.70351 | 0.002359 |
| *pde6a* | XM_039813735.1 | *PDE6A* | 0.701177 | 0.002475 |
| *tmem41aa* | XM_039817756.1 | *TMEM41A* | 0.700037 | 0.002533 |
| *LOC120544521* | XM_039778329.1 | *CIPC* | -0.82163 | 9.48E-05 |
| *slc6a3* | XM_039807297.1 | *SLC6A3* | -0.78493 | 0.000316 |
| *arl16* | XM_039823881.1 | *ARL16* | -0.78248 | 0.00034 |
| *LOC120545153* | XM_039779225.1 | *MEF2C* | -0.759 | 0.000651 |
| *adamtsl5* | XM_039794130.1 | *ADAMTSL5* | -0.75741 | 0.000679 |
| *LOC120544064* | XM_039777595.1 | *TMEM164* | -0.75202 | 0.000779 |
| *pip4p2* | XM_039821087.1 | *PIP4P2* | -0.74411 | 0.000948 |
| *areg* | XM_039802087.1 | *AREG* | -0.74311 | 0.000972 |
| *klhl17* | XM_039801927.1 | *KLHL17* | -0.74149 | 0.001011 |
| *fam120b* | XM_039795670.1 | *FAM120B* | -0.73926 | 0.001067 |
| *dram2b* | XM_039798976.1 | *DRAM2* | -0.72332 | 0.001542 |
| *LOC120560625* | XM_039803303.1 | *CEP120* | -0.72328 | 0.001543 |
| *LOC120559933* | XR_005639388.1 | *NA* | -0.71929 | 0.001686 |
| *LOC120547353* | XR_005637056.1 | *NA* | -0.717 | 0.001772 |
| *fam126a* | XM_039820770.1 | *HYCC1* | -0.71461 | 0.001867 |
| *LOC120550569* | XM_039787169.1 | *TRIM16* | -0.71175 | 0.001985 |
| *map3k14a* | XM_039824035.1 | *MAP3K14* | -0.71164 | 0.001989 |
| *cbx8b* | XM_039816174.1 | *CBX8* | -0.70928 | 0.002092 |
| *trh* | XM_039800554.1 | *TRH* | -0.70785 | 0.002156 |

## G.

| GENE_ID | TRANSCRIPT_ACCESSION_NUMBER | HUMAN ORTHOLOGS | GS | *p-value* |
|---|---|---|---|---|
| *LOC120560720* | XM_039803505.1 | *SEC14L2* | 0.813423 | 0.000127 |
| *si:dkey-6n21.13* | XM_039823385.1 | *P2RY2* | 0.786148 | 0.000305 |
| *gpr176* | XM_039782698.1 | *GPR176* | 0.78102 | 0.000355 |
| *esr1* | XM_039780941.1 | *ESR1* | 0.758584 | 0.000658 |
| *LOC120545314* | XM_039779562.1 | *NA* | 0.748516 | 0.000851 |
| *c20h14orf180* | XM_039785648.1 | *CNST* | 0.744554 | 0.000938 |
| *fbln1* | XM_039808895.1 | *FBLN1* | 0.737168 | 0.001121 |
| *LOC120564637* | XM_039809793.1 | *KBTBD13* | 0.729903 | 0.001328 |
| *zbtb44* | XM_039781008.1 | *ZBTB44* | 0.723127 | 0.001548 |
| *LOC120547035* | XM_039782385.1 | *ESRRG* | 0.722119 | 0.001584 |
| *LOC120572928* | XM_039822453.1 | *BTNL9* | 0.715178 | 0.001844 |
| *irx6a* | XM_039794924.1 | *IRX6* | 0.712514 | 0.001953 |
| *nrros* | XM_039789546.1 | *NRROS* | 0.709855 | 0.002066 |
| *rnf6* | XM_039789936.1 | *RNF6* | 0.70552 | 0.002263 |
| *tacr3a* | XM_039805998.1 | *TACR3* | 0.702018 | 0.002433 |
| *mapk8b* | XM_039787632.1 | *MAPK8* | 0.700128 | 0.002528 |
| *LOC120550569* | XM_039787169.1 | *TRIM16* | -0.8027 | 0.000182 |
| *p2ry2.1* | XM_039822179.1 | *P2RY2* | -0.76879 | 0.000501 |
| *LOC120559933* | XR_005639388.1 | *NA* | -0.76254 | 0.000593 |
| *LOC120567767* | XM_039814778.1 | *CCL25* | -0.75263 | 0.000767 |
| *nmur3* | XM_039796436.1 | *NMUR2* | -0.73464 | 0.00119 |
| *ogfod2* | XM_039802729.1 | *OGFOD2* | -0.72915 | 0.001351 |
| *anapc10* | XM_039823076.1 | *ANAPC10* | -0.72902 | 0.001355 |
| *LOC120563179* | XM_039807335.1 | *NPR1* | -0.72414 | 0.001514 |
| *prr15lb* | XM_039789397.1 | *PRR15L* | -0.71812 | 0.00173 |
| *scn1lab* | XM_039793357.1 | *SCN2A* | -0.71675 | 0.001782 |
| *LOC120569990* | XM_039818213.1 | *CYBRD1* | -0.70614 | 0.002234 |
| *LOC120569118* | XM_039816997.1 | *THAP2* | -0.70318 | 0.002375 |
