## Supplementary file 4 for "Does transcriptome of freshly hatched fish larvae describe past or predict future developmental trajectory?"

**Supplementary file S4:** Primers used for validation and normalization of expression level of candidate genes

| | Primers | |
|---|---|---|
| Gene symbol | **Forward** | **Reverse** |
| *selenoo* | CGTTGGGCTCTGAACTGGAT | TCCGGCTCTTCTTTTCCCAC |
| *nudt12* | TCCTGCTTGGTTTGCCATCA | GCCTCTTCCTCGTTGAGCTT |
| *sec14l2* | CTTGGGAGGAACGTGGAGTC | CAGGAAAGAGTTTGGGGGCT |
| *trim16* | AAAGCCTGTCCTCCCTCCTG | CTCTGCTCCATGTGTTGCCT |
| *mkx* | TACAAGCACCGAGACAACCC | TGCCTCACCGTGTTCTTCAG |
| *crcp* | TTCCTGACCACCATGATGCC | TGCTCCTCTGATAACCGCTC |
| *atp8a2* | GCCAGATCCTTTTTGAGCGC | AGCATTCTGGGTGATTCGGT |
| *slc15a1* | TGACAACGATCCTCTGGTGC | GTGTTTGTCGCCGTCCTTTG |
| *prdm1* | AAACAGCACCTACCTCAGCA | ATCTGGTCGTCGGGTGTGTC |
| *txnl4a* | GGTGGACATCACAGAAGTGC | TCTCCTGCTTGTCCTCCATT |
| *aoc1* | CCAAACCAACCCCAACATCAC | GCCTTGACTCAAAGCGGACA |
| *cipc* | TAAAGAGAGGACACAGCCGC | CCTTTCCCTCATGGCTCTCC |
| *pycard* | TCGCAGTCATTTTGGATGAGC | GCTTTCAGAGGCCCAGAGTAG |
| *si:dkey-117m1.4* | TCATCACAAACTGCGGGACA | GCGGGCTTCTCTTTAGGACA |
| *Houskeeping genes* | | |
| *txn2* | CGCGAGGTCTCCTTTAACGT | ATCGCCAGGTCTGTGTGATC |
| *acadl* | ATCTTCAGGCAAAGCGTCCG | ATACATTTGCTCCTCCCACG |
| *gsta.1* | AGCAAAGGACCGCTACCTTC | CCTCCAACATCAGGGTGCAT |
| *wdr83os* | TGAAGTGGTGTGCCTGGATC | GCTGTGGGTTCTGGAGGTAC |

.
