## Supplementary file 5 for "Does transcriptome of freshly hatched fish larvae describe past or predict future developmental trajectory?"

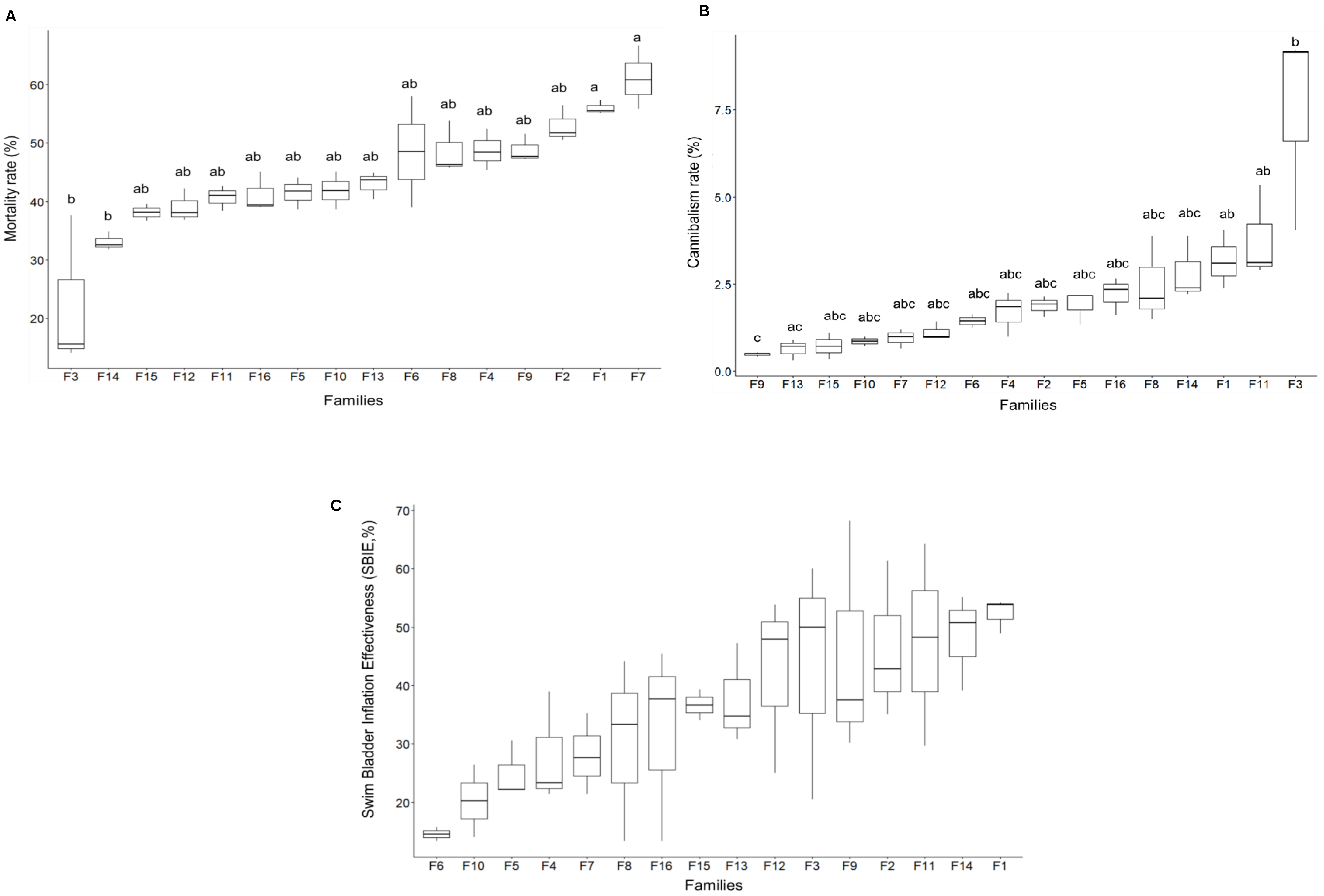


**Fig.S1: A: Box plot diagrams of mortality rate**. **B. Box plot diagram of Cannibalism rate**. **C. Box plot diagram of swim bladder inflation effectiveness rate (SBIE).** The graphs show statistical comparisons between the lengths of larvae from 16 E. perch families at different developmental stages. The experimental groups are arranged in an ascending order according to their mean values and the letters indicate significant differences ($p<0.05$) between the groups.

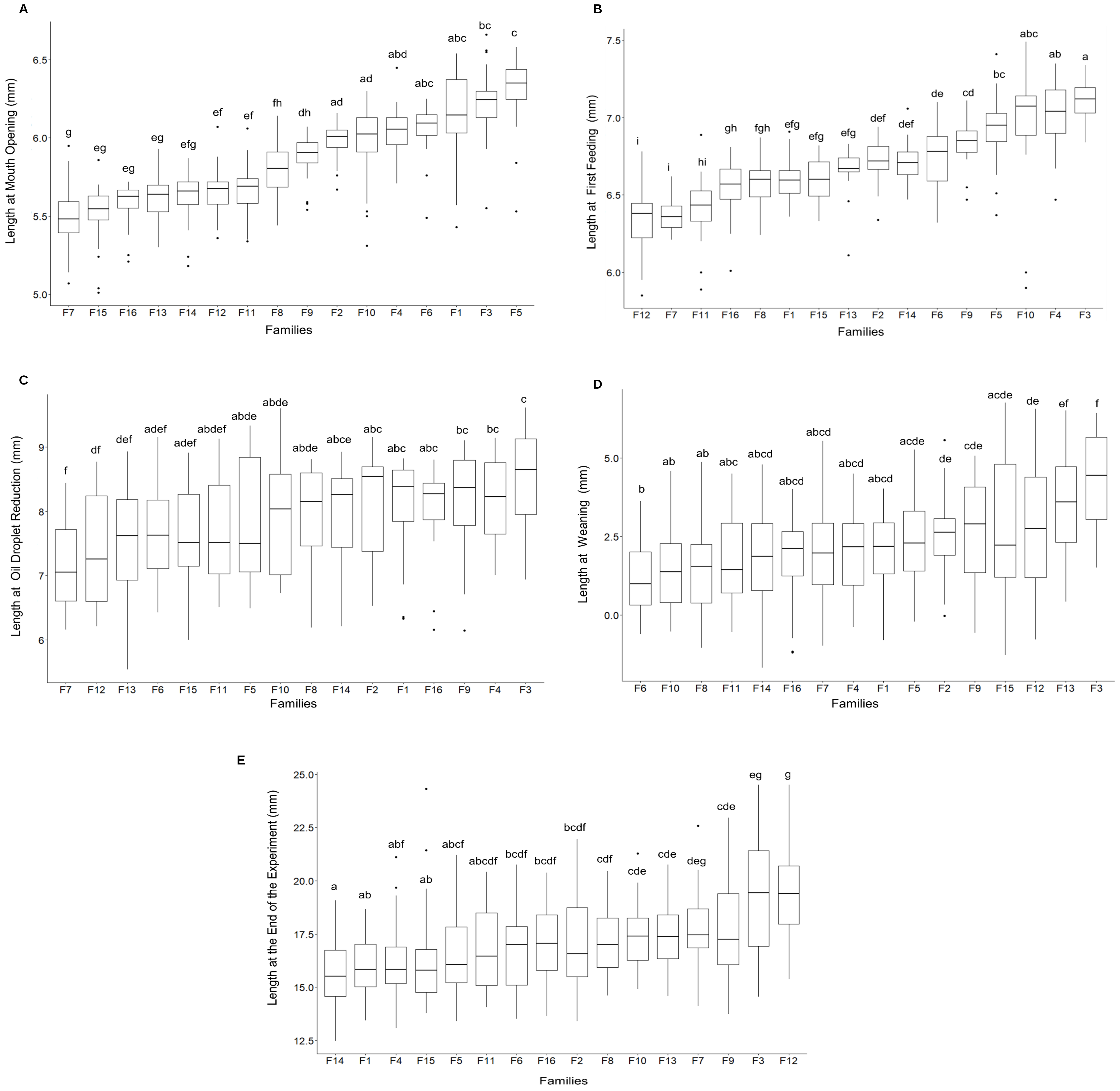

**Fig.S2:** Box plot diagrams of total length of fish larvae at **A.** mouth opening. **B.** first feeding stage. **C**. oil droplet reduction stage. **D**. weaning stage. **E**. the end of the experiment. The graphs show statistical comparisons between the lengths of larvae from 16 E. perch families at different developmental stages. The experimental groups are arranged in an ascending order according to their mean values and the letters indicate significant differences (p<0.05) between the groups.

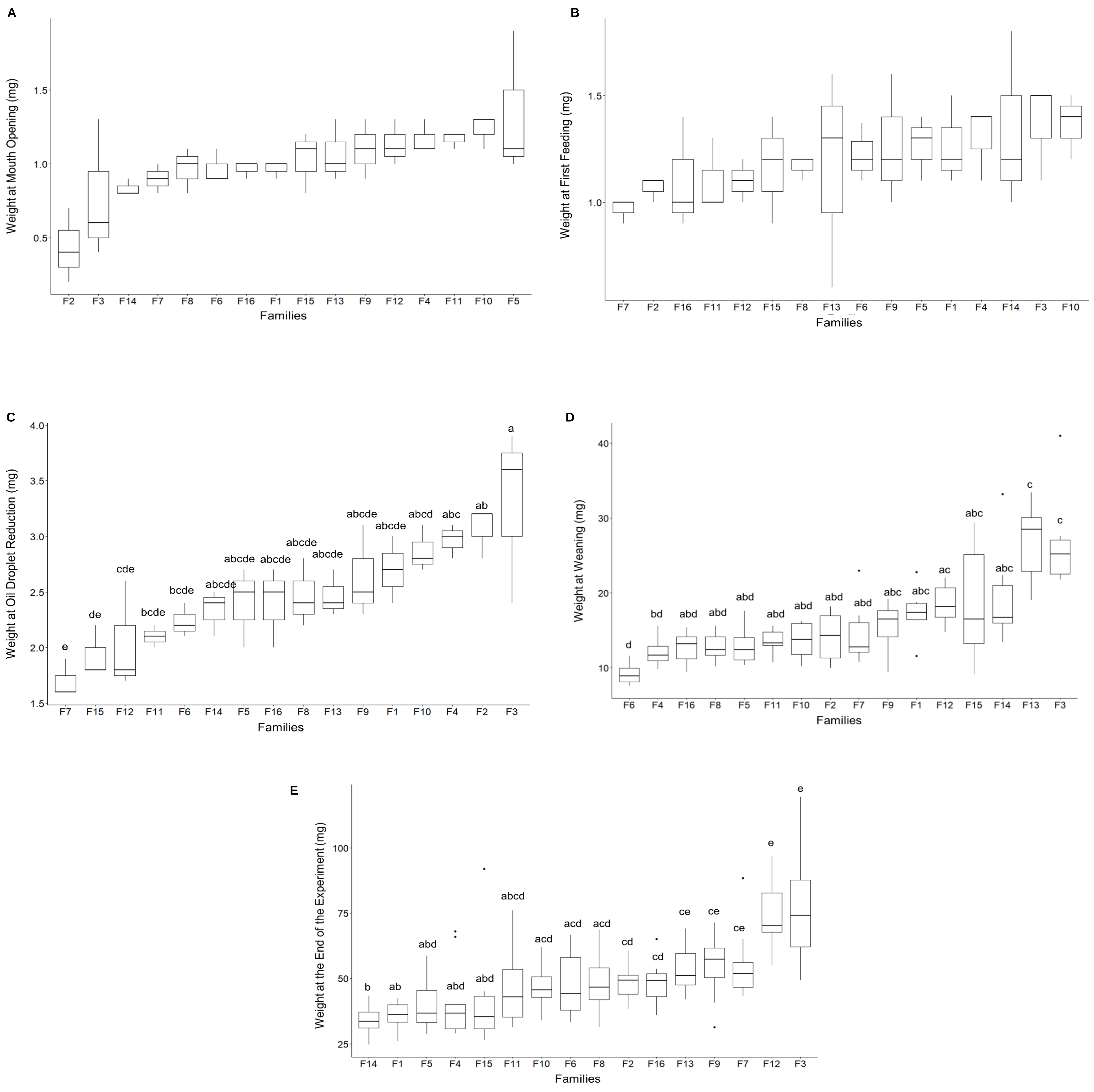


**Fig.S3:** Box plot diagrams of weight of fish larvae at **A.** mouth opening. **B.** first feeding stage. **C.** oil droplet reduction stage. **D.** weaning stage. **E.** the end of the experiment. The graphs show statistical comparisons between the weights of larvae from 16 E. perch families at different developmental stages. The experimental groups are arranged in an ascending order according to their mean values and the letters (when present) indicate significant differences (p<0.05) between the groups.

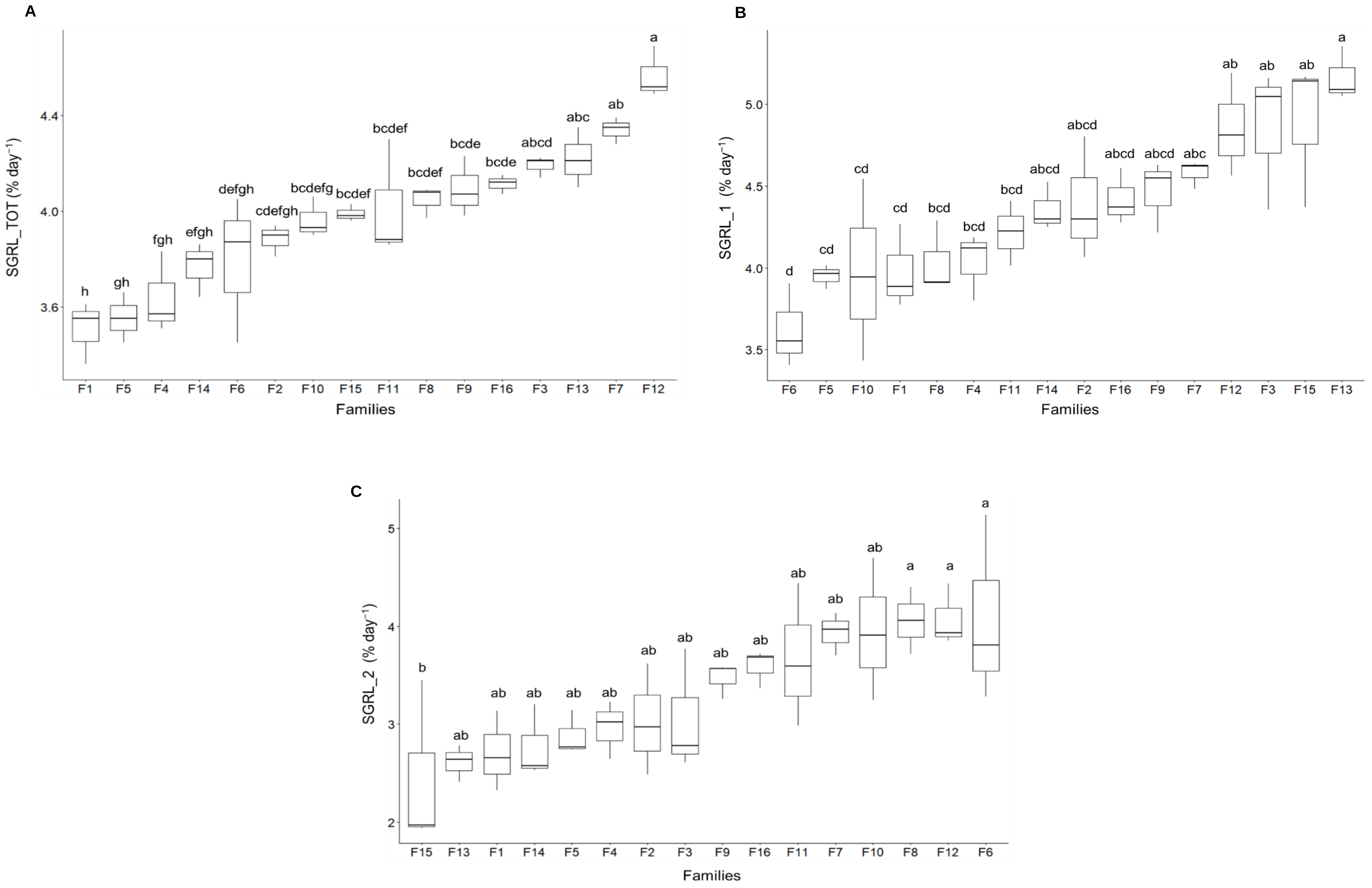


**Fig.S4:** Box plot diagram of **A.** Specific growth rate for length (SGRL_TOT) of larvae considering the entire rearing period (0 DPH-27DPH). **B**. Specific growth rate for length (SGRL_1) of larvae from hatching until the weaning stage (0 DPH-17DPH). **C.** Specific growth rate for length (SGRL_2) of larvae from weaning until the end of the experiment (17 DPH- 27 DPH). The graphs show statistical comparisons between the SGR of larvae from 16 E. perch families at different developmental stages. The experimental groups are arranged in an ascending order according to their mean values and the letters indicate significant differences ($p<0.05$) between the groups.

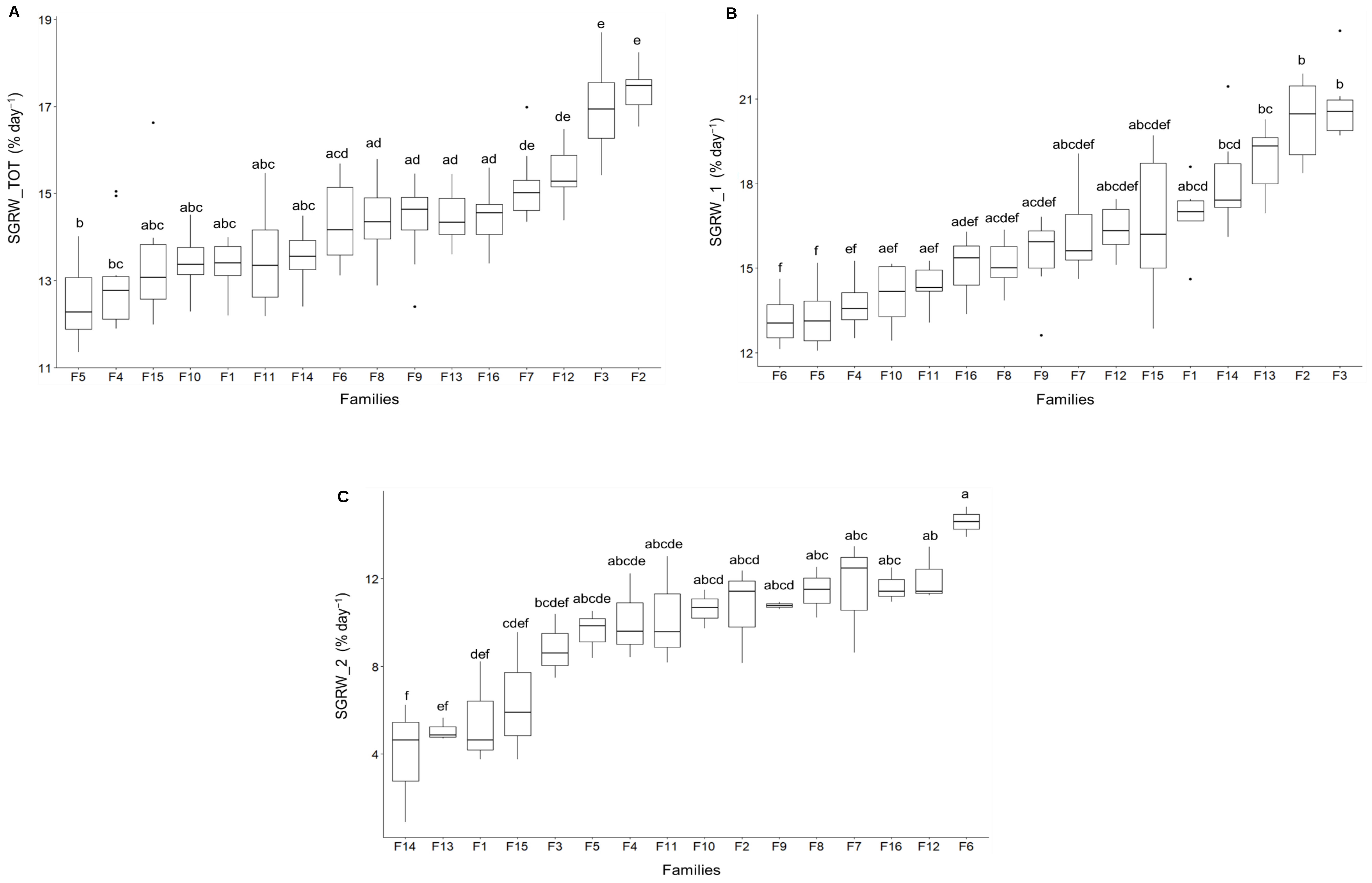


**Fig.S5:** Box plot diagram of **A.** Specific growth rate for weight (SGRW_TOT) of larvae considering the entire rearing period (0 DPH-27DPH). **B**. Specific growth rate for weight (SGRW_1) of larvae from hatching until the weaning stage (0 DPH-17DPH). **C.** Specific growth rate for weight (SGRW_2) of larvae from weaning until the end of the experiment (17 DPH- 27 DPH). The graphs show statistical comparisons between the SGR of larvae from 16 E. perch families at different developmental stages. The experimental groups are arranged in an ascending order according to their mean values and the letters indicate significant differences ($p<0.05$) between the groups.

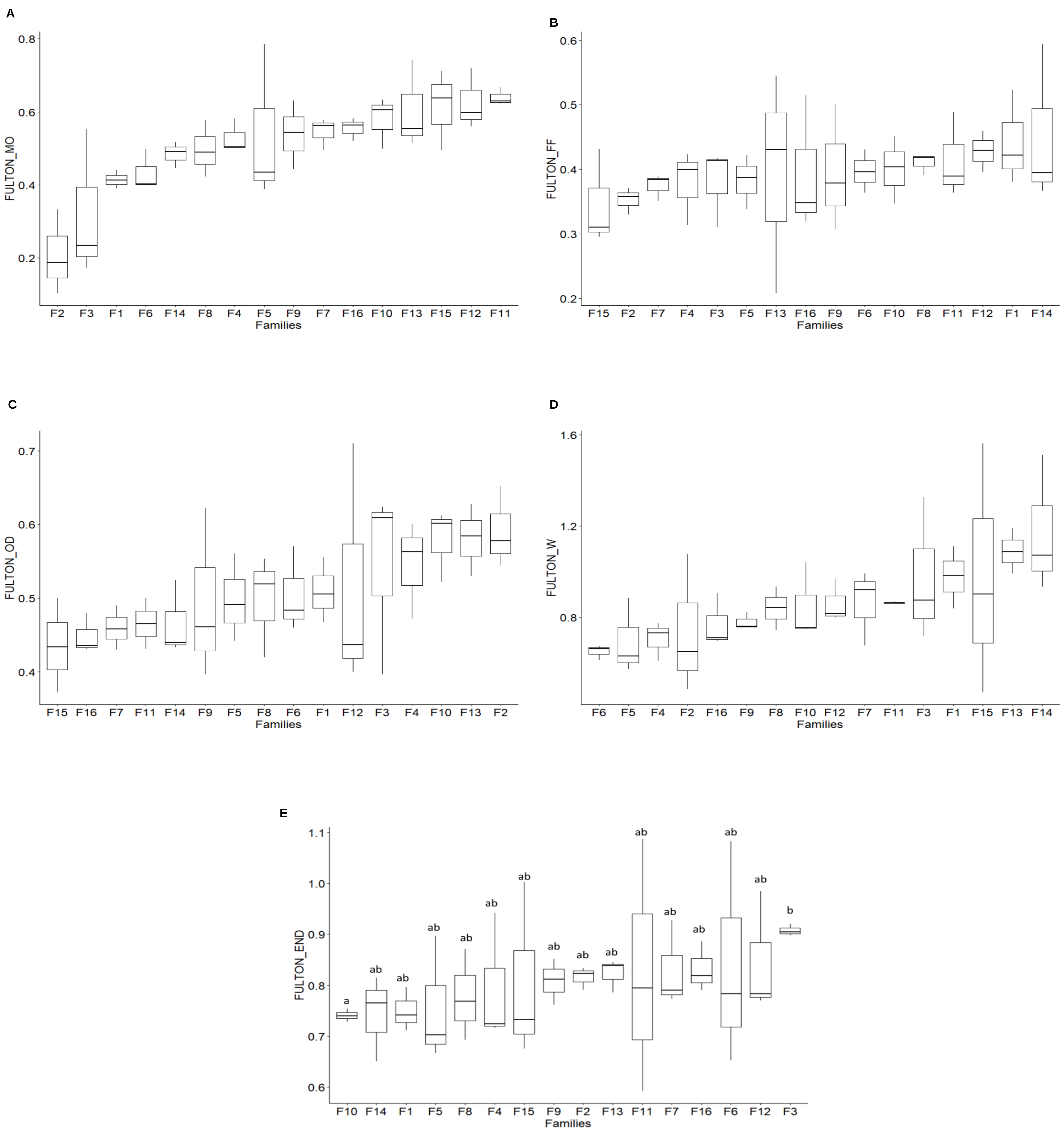


**Fig.S6:** Box plot diagram of A. Fulton's condition factor (K) of larvae at mouth opening. **B**. Fulton's condition factor (K) of larvae at first feeding stage. **C**. Fulton's condition factor (K) of larvae at oil droplet reduction stage. **D**. Fulton's condition factor (K) of larvae at Weaning stage. **E**. Fulton's condition factor (K) of larvae at the end of the experiment. The graphs show statistical comparisons between the K of larvae from 16 E. perch families at different developmental stages. The experimental groups are arranged in an ascending order according to their mean values and the letters (when present) indicate significant differences ($p<0.05$) between the groups.

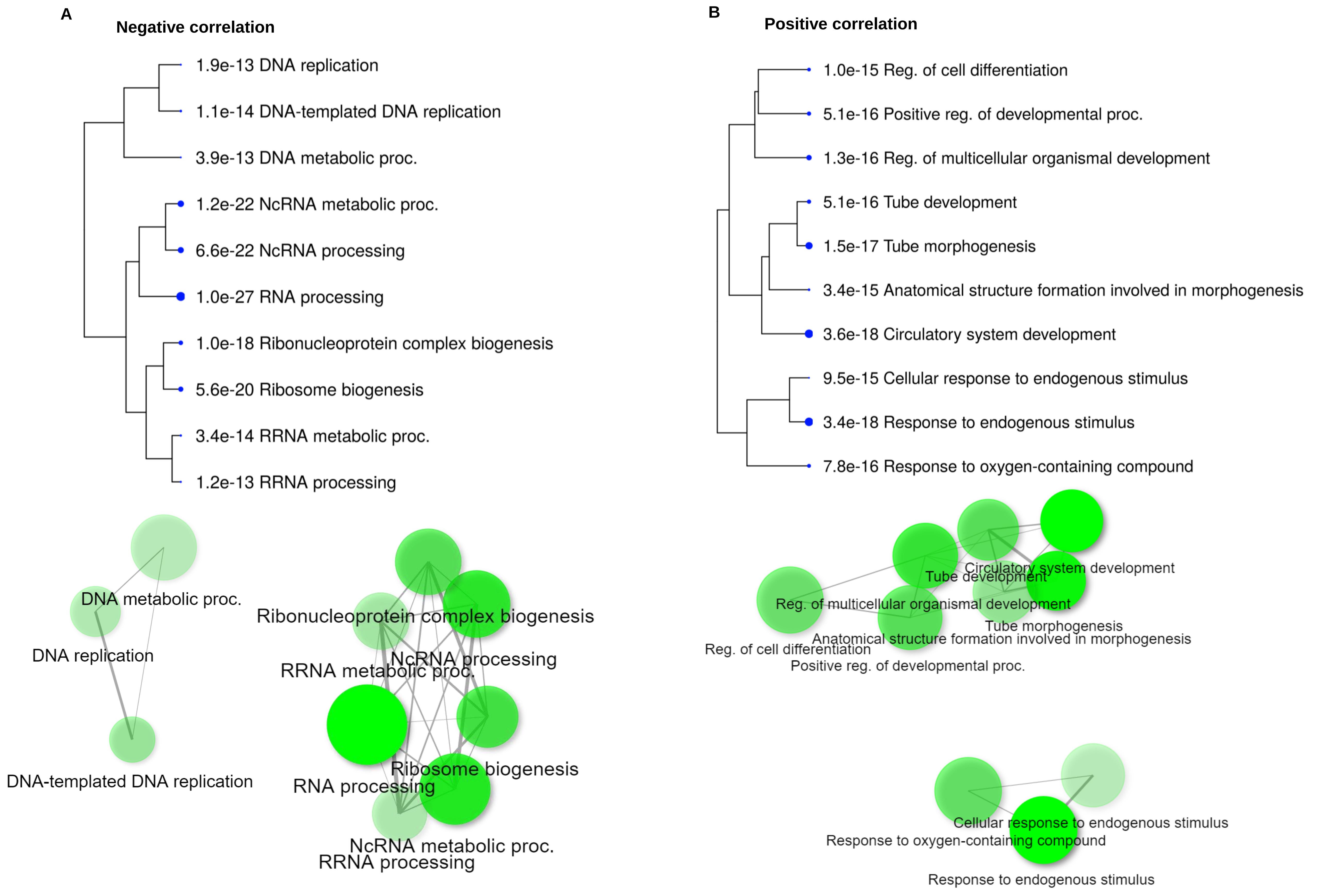


**Fig. S7**: **Tree view and network visualization showing the 10 most significantly enriched GO (biological process)** for most significant gene modules related to embryonic developmental rate**.** A. Negative correlated modules B. Positive correlated modules

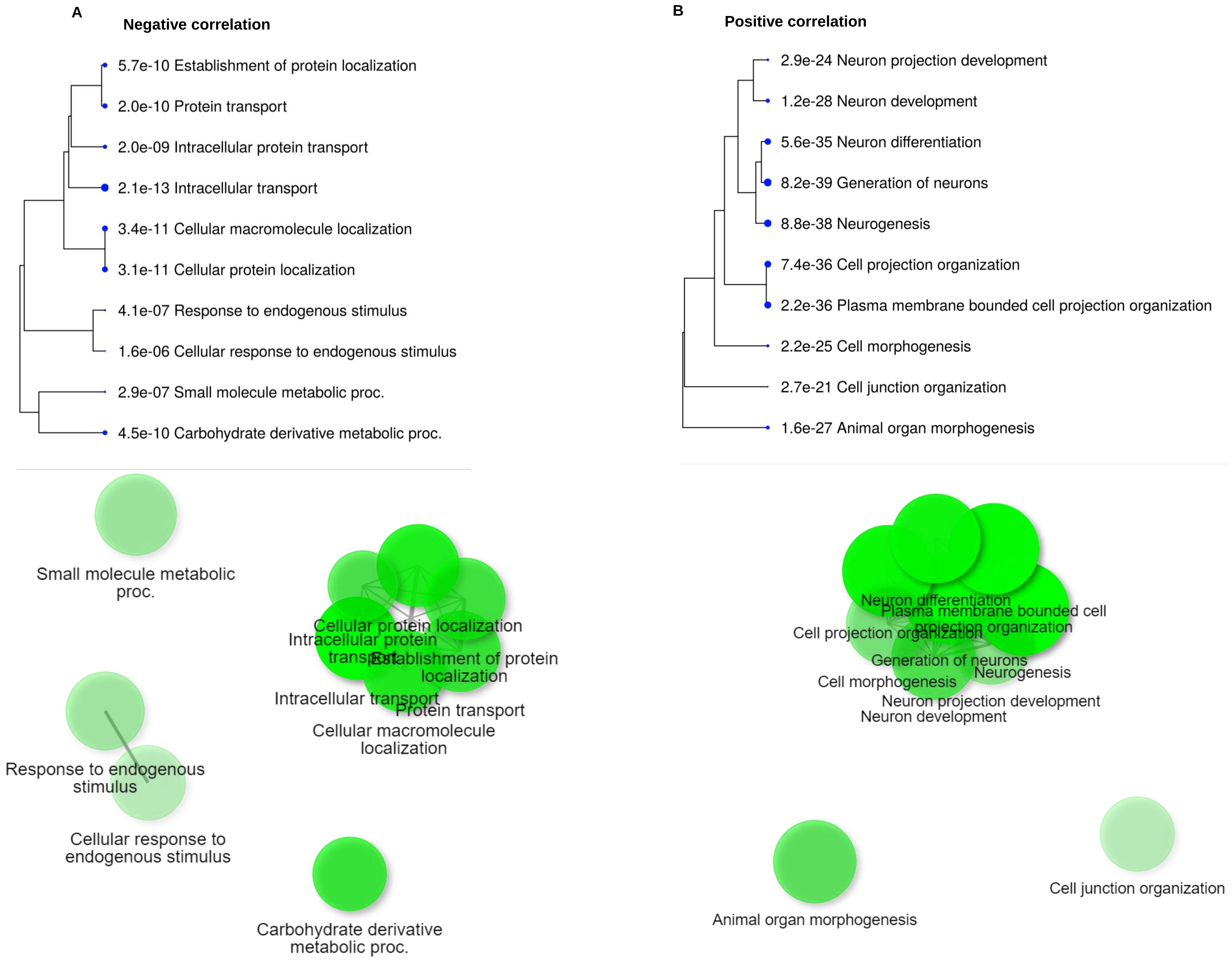


**Fig. S8**: **Tree view and network visualization showing the 10 most significantly enriched GO (biological process)** for the most significant gene modules related to hatching rate. A. Negative correlated modules B. Positive correlated modules

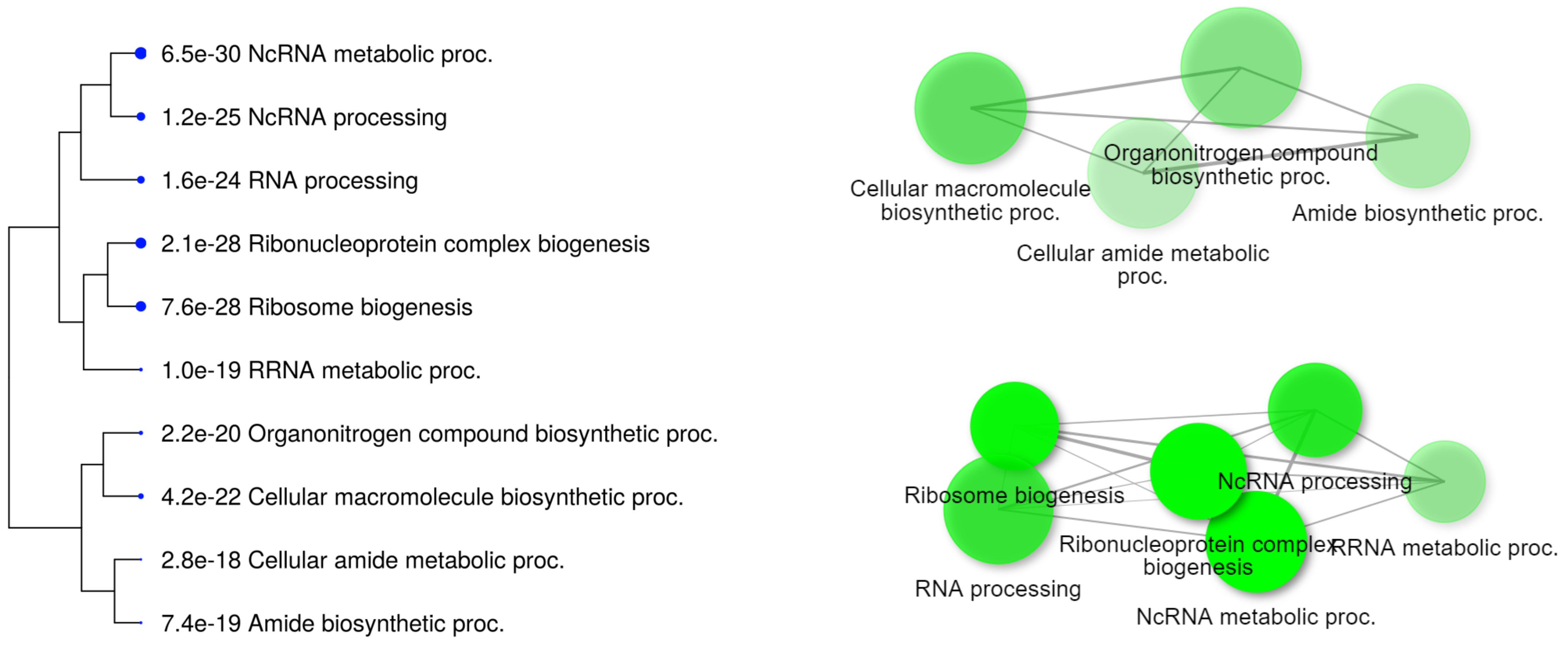


**Fig. S9**: **Tree view and network visualization showing the 10 most significantly enriched GO (biological process)** for most significant gene modules positively related to weight of larvae at mouth opening**.**

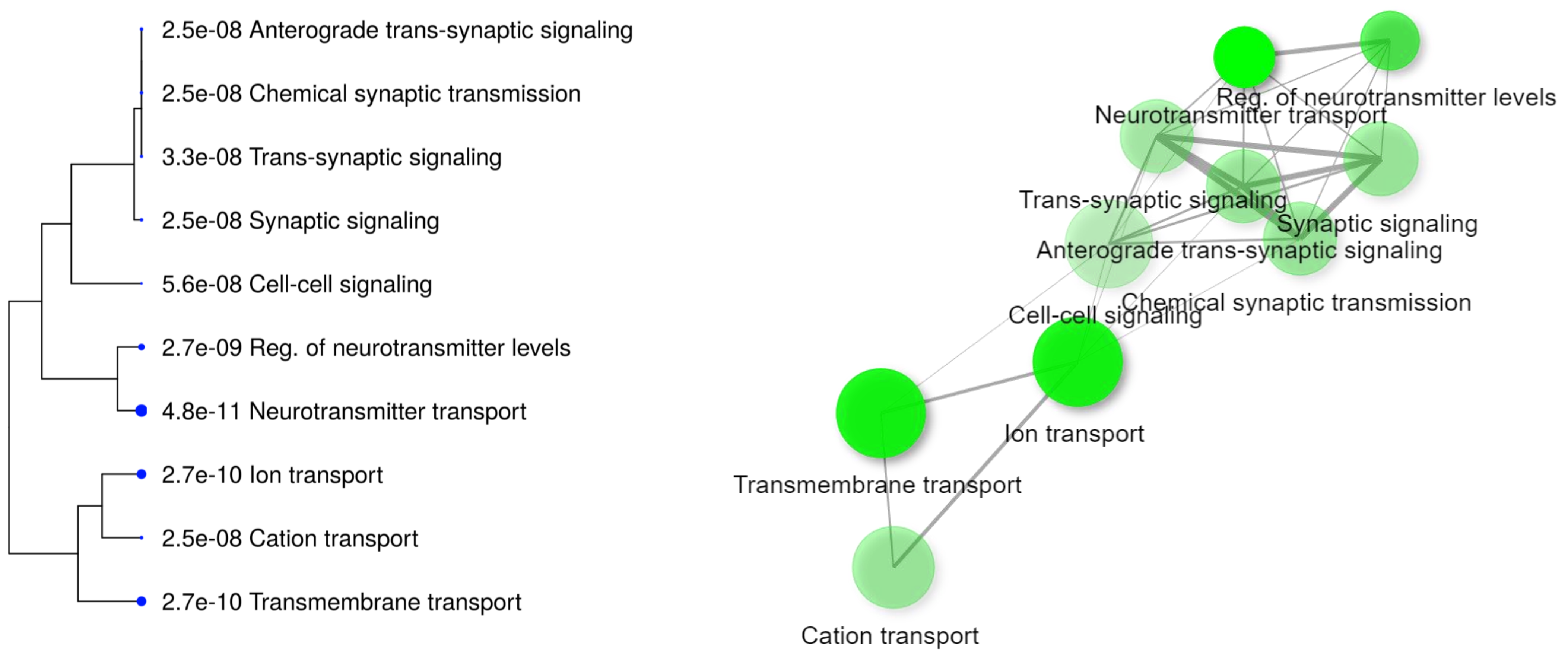


**Fig. S10**: **Tree view and network visualization showing the 10 most significantly enriched GO (biological process)** for most significant gene modules negatively related to length of larvae at first feeding.

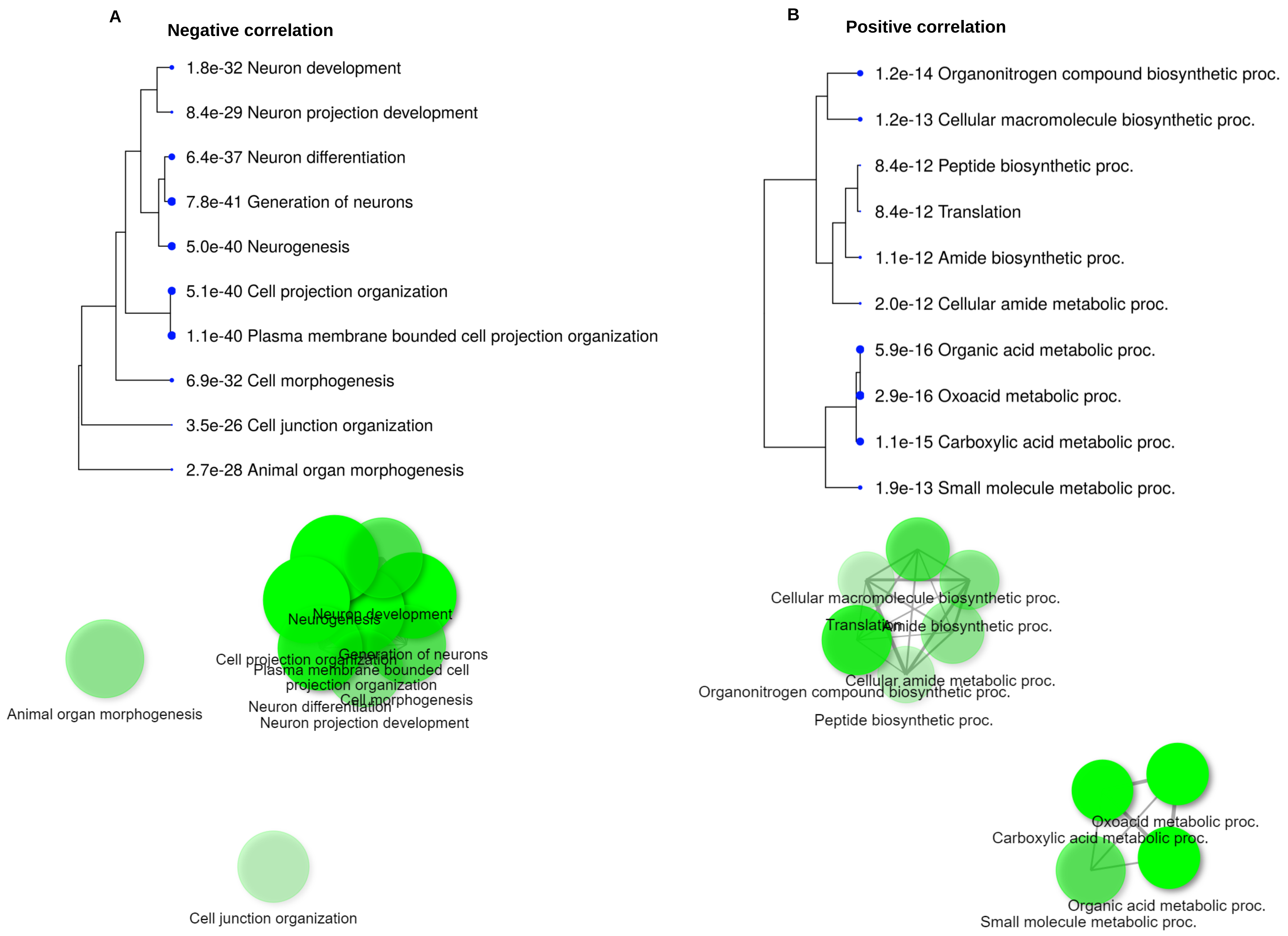


**Fig. S11**: **Tree view and network visualization showing the 10 most significantly enriched GO (biological process)** for most significant gene modules related to Fulton's condition factor (K) of larvae at mouth opening. A. Negative correlated modules. B. Positive correlated modules

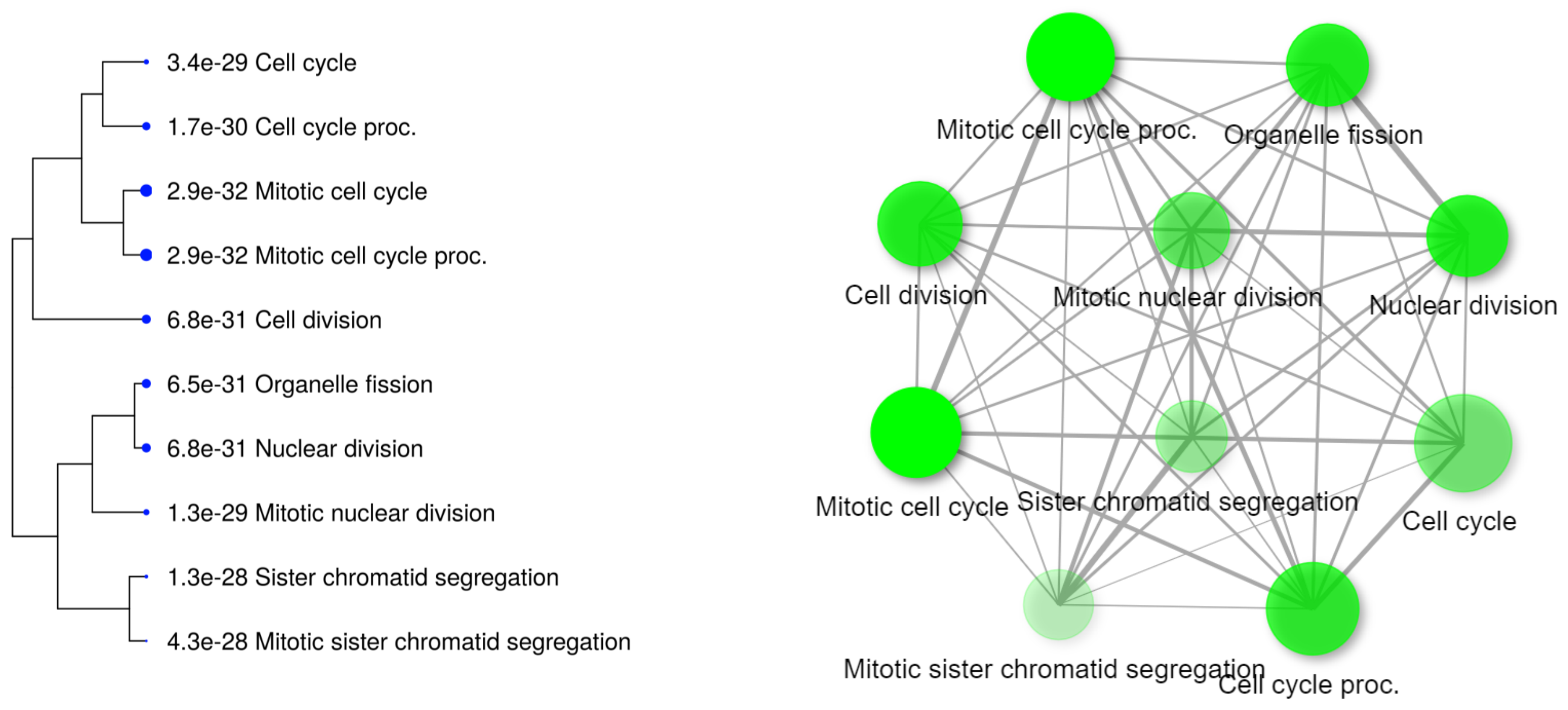


**Fig. S12**: **Tree view and network visualization showing the 10 most significantly enriched GO (biological process)** for most significant gene modules negatively correlated to Fulton's condition factor (K) of larvae at oil droplet reduction stage.

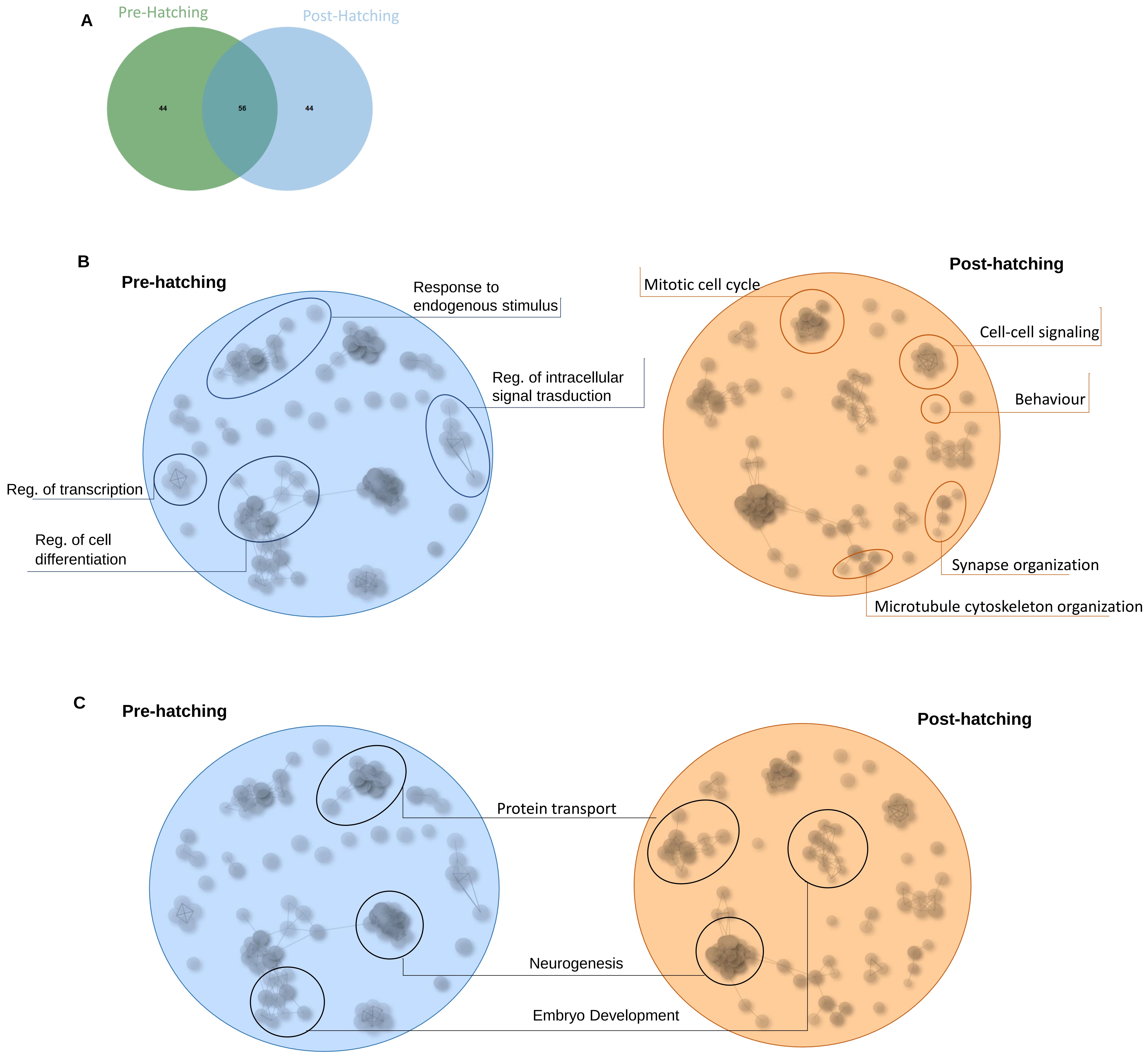


**Fig. S13**: Clustering of the 100 most enriched biological processes (BPs) obtained during the functional enrichment analysis (FDR < 0.05) A. Venn diagram showing the number of pre- and post-hatching specific and common GO terms. **B.** Circled clusters are those indicating pre-hatching and post-hatching traits specific cluster. **C.** Circled clusters are those indicating common clusters between pre- and post-hatching indicators.

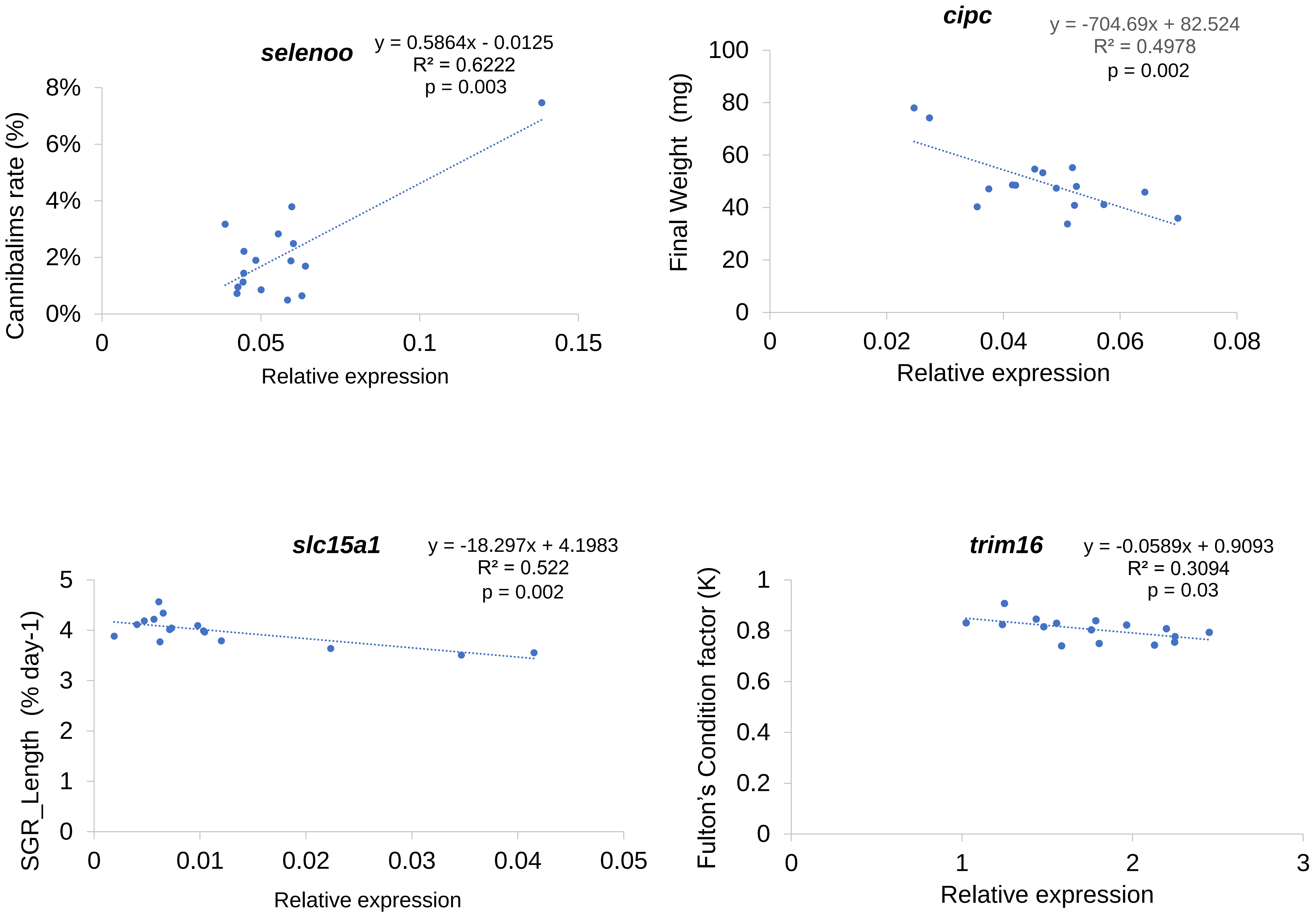


**Fig. S14**: Graphs showing the correlation between qPCR-validated genes and their associated traits
